# *Bacillus* competence pili are efficient single - and double stranded DNA uptake machines

**DOI:** 10.1101/2025.11.24.690153

**Authors:** Alexandra Kilb, Felix Dempwolff, Peter L. Graumann

## Abstract

Natural competence describes the ability of diverse bacterial species to take up DNA from the environment for integration into the host chromosome, allowing the acquisition of novel genetic information. Uptake in many species operates via a dynamic pilus structure that pulls DNA through the cell wall into the periplasm by pilus retraction, followed by transport of one of the two DNA strands into the cytosol by an active membrane transporter. How double stranded DNA with a bending contour of hundreds of base pairs can fit through a narrow pore accommodating a pilus has been a puzzling question. We show that purified major and minor pilins from two *Bacillus* species, *B. subtilis* and *Geobacillus thermodenitrificans*, show efficient binding to double as well as to single stranded DNA with similar affinities. Consistent with uptake, ssDNA accumulates and moves within the periplasm before transport through the cell membrane, with similar dynamics as dsDNA. Thus, *Bacillus* competence pili are dsDNA as well as ssDNA uptake machines. The binding of pili to ssDNA regions within environmental DNA likely increases the efficiency of competent bacteria to pull extended DNA strands through the cell wall. Uptake of environmental ssDNA may also provide a beneficial source of nutrition for bacteria during stationary growth phase.

## Introduction

Natural competence is a major driver of evolution, allowing uptake of large fragments of DNA that may contain novel genetic information (1–4). This gain of genetic traits is also a threat to human health as many human pathogenic bacteria are naturally competent, allowing them to efficiently gain access to resistance genes. Competence has so far been considered to apply to the uptake of double stranded DNA from the environment, which has to cross the outer membrane and cell wall of Gram-negative or the thick cell wall of Gram-positive bacteria (4). How type IV pili, which have been recognized to be the major machinery used by competent bacteria for DNA uptake through the cell wall, pull a large molecule like DNA that cannot be easily bent through the tight peptidoglycan meshwork of the cell wall, is still puzzling.

Surface appendages like type IV pili play a crucial role for bacteria in functions such as twitching motility, surface attachment, biofilm formation, or uptake of environmental DNA by natural transformation (5). In the 1970s, type IV pili were first described for *Pseudomonas aeruginosa* and since then have been identified in a vast number of bacteria including many human pathogens (6). Although the functions of these appendages are diverse, their overall architecture is largely conserved (7,8). Pilus assemblies in Gram-negative bacteria tend to be more complex than in Gram-positives mainly due the presence of components forming a ring within the outer membrane (9). Type-IV pilins (T4P) are divided into two groups, T4aP and T4bP (10) dependent on their sequence, length of the leader peptides as well as differences in their assembly (11). T4aP are involved in twitching motility and also DNA uptake (12), while T4bP play a crucial role in biofilm formation, bacterial colonization and cell adhesion (13). All T4P share a common structural setup, where a helical filament is formed by the assembly of thousands of copies of pilins (11). Prior to assembly into the pilus structure, the synthesized pre-pilin molecules have to mature into the functional pilin unit by cleavage of the signal peptide, which is removed by a prepilin-peptidase upon membrane integration via the Sec machinery (14,15). This cleavage is necessary for the polymerization of the pilins (16). T4P are thought to be largely composed of so called major pilins, while minor pilins assemble at the tip of the polymer in some bacteria (17,18), where they have been shown to bind to DNA (19). Alternatively, they can be incorporated into the entire pilus structure (20). Assembly into a filament structure depends on an assembly ATPase and an inner membrane core protein in all bacteria analysed so far (21–25). A model has been proposed that the assembly ATPase undergoes a conformational change upon binding of ATP, which is further transmitted through the core protein, functioning as a “lever” whereby the pilus is pushed out of the membrane; this might allow new pilin monomers to click into a gap at the base of the pilus filament (26), leading to filament extension. Some bacteria possess an additional ATPase mediating pilus retraction, which may be achieved by the assembly ATPase in species missing the retraction ATPase (8). Alternation between extension, adhesion and retraction may promote the transport of environmental DNA into the periplasm (2,27,28).

Competence can be induced by antibiotics or DNA-damaging agents, such as in *Streptococcus pneumoniae* (29), or by population density control (quorum sensing) due to peptide pheromones, so called competence-stimulating-peptides (CSPs) (30). In the model organism *Bacillus subtilis,* natural competence is induced only in a subpopulation during entry into stationary growth phase. CSPs stimulate the stabilization of master transcription factor ComK, which in turn mediates upregulation of about 100 genes (10,31–33). The components of the cell envelope-spanning machinery for DNA uptake are mainly encoded in the *comG* operon, containing all core proteins of type IV pilus filaments: hexameric ATPase ComGA, putative platform protein ComGB, major pilin ComGC and several minor pilins (ComGD-ComGG) (34,35). Double stranded DNA (dsDNA) is taken up by the retraction of the pilus structure, and will be bound by the membrane-bound DNA receptor ComEA in the periplasm. ComEA binds single (ss-) as well as dsDNA *in vivo* and *in vitro*, with a higher affinity for dsDNA (36,37). One strand of the incoming (transforming) DNA is degraded, while the other one passes through the membrane channel ComEC, with the ATPase ComFA likely powering the transport. Inside the cytosol, RecA guides ssDNA onto the chromosome where it can be integrated if sufficient homology exists (38–40).

While membrane transport of incoming DNA and chromosome integration are relatively well understood, the initial process of DNA uptake via binding to the competence pilus is poorly characterized. *T. thermophilus* or *V. cholerae* appear to bind DNA at the tips of pili via minor pilins (19,41) while *S. pneumoniae* pili appear to bind DNA at their entire length (42). To fit through a narrow pore within the cell wall accommodating a pilus, dsDNA would have to be bound at its ends, but the more likely case of binding to dsDNA fragments internally leads to the problem that dsDNA having a persistence length of hundreds of base pairs cannot be bent to fit into a pore-like structure.

We have studied DNA binding properties of major and minor pilins from two *Bacillus* species, thriving in different habitats. We show that *B. subtilis* major pilin ComGC binds to dsDNA *in vitro,* and similarly major pilin ComGC from *Geobacillus thermodenitrificans* and also its minor pilin ComGD. Thus, both classes of pilins have DNA-binding capacity. Interestingly, we also find binding of major and minor pilins to single stranded DNA, even at high salt concentration. ComGD shows similar affinity to ss - and dsDNA in the lower micromolar range. Because we find that ssDNA fragments are efficiently transported into the periplasm of *B. subtilis*, our data provide evidence that *Bacillus* pili can efficiently bind to and transport ss - and dsDNA.

Binding of pili to ssDNA regions within environmental DNA may increases the efficiency of competent bacteria to pull extended DNA strands through the cell wall, and allows cells to access environmental ssDNA as nutrient source.

## Materials and methods

### Growth conditions

*Escherichia coli* cells were grown in lysogeny broth (LB) medium and on 1.5 % agar plates supplemented with 100 µg/ml ampicillin [amp] at 37°C. If cells were used for protein overexpression, a transformation of *E. coli* cells with the respective plasmid was performed the day prior the experiment. For inoculation, cells were scratched from a plate and inoculated in LB-media supplemented with 1.2% lactose for heterologous protein expression. For all constructs, cells were incubated over night at 200 rpm at 30°C. In case of *B. subtilis*, cells were grown on 1.5% LB-agar plates and in LB-medium at 37°C, supplemented with the respective antibiotics (if necessary) to the following concentrations: 25 μg/ml lincomycin and 1 μg/ml and erythromycin [MLS], and 25 μg/ml kanamycin [kan]. To induce natural competence in *B. subtilis* cells for transformation efficiency assays or microscopy experiments, a modified competence medium was used according to the method of J. Spizizen (100 ml 10 x MC-media composed: 14.01 g K_2_HPO_4_ × 3 H_2_O, 5.24 g KH_2_PO_4_, 20 g glucose, 10 mL trisodium citrate (300 mM), 1 ml ferric ammonium citrate (22 mg/ml), 1 g casein hydrolysate, and 2 g potassium glutamate (sterile filtered)) (43). For inoculation of a 10 ml culture, 1 ml of 10x MC media, 0.333 ml of sterile 1 M MgSO_4_, the respective antibiotics or supplements were added and adjusted with ddH_2_O to a final volume of 10 ml. Cells were grown to competence on a shaking platform (200 rpm) at 37°C. Strains used in this study are listed in table S5.

### Strain constructions

Oligonucleotides and generated plasmids are listed in table S6 and S7. PCR fragments were generated by using chromosomal DNA of PY79 wild-type, *Geobacillus thermodenitrificans* wild-type cells or vector pDG1164 (depending on the construct). The genes encoding for respective proteins were amplified without their N-terminal transmembrane domain for protein overexpression. To generate recombinant plasmids, modular cloning was used with the restriction enzyme *Bsa*I and the T4-DNA-ligase (New England Biolabs). For this, the vector and the insert were digested with *Bsa*I at 37°C for 4 minutes, followed by a ligation step at 16°C for 5 minutes with 5 repetitions and a final ligation step at 16°C for 10 minutes. DH5α cells were transformed with the resulting plasmid and plasmids were checked by Sanger sequencing by using a T7 promotor regions. In case of *comGC* from *B. subtilis*, construct was cloned into a His-GST containing vector, *comGC* as well as *comGD* from *G. thermodenitificans* in an His-MBP containing vector, resulting in N-terminal fusion strains. Each vectore encodes a TEV-protease recognition site (ENLYFQS) distal to the tag and proximal to the protein of interest. To achieve single base pair substitutions to express point mutant variants, Q5 directed mutagenesis was used. Primers were designed with the NEB primer design tool (New England Biolabs) and prior generated plasmids were used as templates. Mutagenesis was done according to manufactures instructions (New England Biolabs) (table S1). Genome editing of *B. subtilis* to generate point mutations was done by CRISPR-cas9 according to Burby and Simmons 2017 (44).

### Protein purification

For protein purification *E. coli* C41 cells were transformed with plasmids the day prior the experiment and six liters of LB-media were incubated with cells over night at 30°C on a shaking platform (120 rpm) with 1.2% lactose for protein overexpression (for all used constructs). Cells were harvested at 4000 rpm for 15 minutes (Beckmann centrifuge, JLA 9.1000 rotor) and the generated cell pellet was resuspended in 10 ml of lysis buffer per gram of cells. Depending on the construct, different buffer compositions were used: His-GST-ComGC (20 mM Hepes, 1 M NaCl, 40 mM imidazole, 20 mM KCl, 20 mM MgCl_2_); His-MBP-ComGC (20 mM Hepes, 1 M NaCl, 40 mM imidazole, 20 mM KCl, 20 mM MgCl_2,_ 1 mM DTT, 10% glycerol); His-MBP-ComGD (20 mM Hepes, 250 mM NaCl, 40 mM imidazole, 20 mM KCl, 20 mM MgCl_2_). To disrupt the cells, a microfluidizer (microfluidics) was used and cell debris was removed by centrifugation at 15000 rpm for 20 minutes using a Beckmann rotor JA-25.50. Filtered lysate (0.45 µm, Filtropur) was subsequently applied to a 5 ml His-trap column (GE healthcare) which was further pre-equilibrated with the respective buffer. After applying the sample to the column, a washing step (with at least 3 CV) followed before the elution with elution buffer for His-GST-ComGC (20 mM Hepes, 1 M NaCl, 250 mM imidazole, 20 mM KCl, 20 mM MgCl_2_); His-MBP-ComGC (20 mM Hepes, 1 M NaCl, 250 mM imidazole, 20 mM KCl, 20 mM MgCl_2_1mM DTT, 10% glycerol); His-MBP-ComGD (20 mM Hepes, 250 mM NaCl, 250 mM imidazole, 20 mM KCl, 20 mM MgCl_2_). To concentrate the protein a Amicon Ultracel-3K or Amicon Ultracel-10K (Millipore) was used, depending on the size of the protein.

In case of His-GST-ComGC the affinity tag was removed by treatment with TEV-protease overnight at room temperature. TEV-protease (which possessed a His-tag itself) and the affinity-tag (His-GST) were separated from ComGC via reverse-nickel affinity chromatography by a 5 ml His-trap column. For this, the column was equilibrated with Buffer (20 mM Hepes, 1 M NaCl, 40 mM Imidazole, 20 mM KCl, 20 mM MgCl_2_). Flow through, washing steps were collected separately in three distinct fractions of 15 ml and examined for presence of protein by SDS PAGE containing 1 mM DTT as reducing agent. Fractions including the protein were concentrated with an Amicon Ultracel-3K.

### Preparative and analytical size exclusion chromatography

Size exclusion chromatography was used to remove impurities and aggregated proteins from elution fraction. For this we used an NGC-system (Bio-Rad) with a Superdex XK16/600 column (Cytivia, Massachusetts, USA). Before use, the column was equilibrated with a 200 mM HEPES buffer (pH 7.5, “SEC buffer”), in case of His-MBP-*Gt*ComGC 0.75 M NaCl, 10% glycerol and 1 mM DTT was added to the buffer in case of *Bs*ComGC 0.75 M NaCl. For calibration a standard protein mix was used with known molecular weights (Thyroglobulin (bovine) 670 kDa; γ-Globulin (bovine) 15 kDa 8; Ovalbumin (chicken) 44; Myoglobin (horse) 17; Vitamin B12 1.35 kDa) (Biorad). For analytical size exclusion chromatography the same NGC-system was used with a Superdex 200 Increase 10/300 column (Cytivia). Before use, column was equilibrated with the respective buffer. Chromatograms were generated with the NGC software and Microsoft Excel 2019.

### Electromobility shift assay (EMSA)

To determine if the recombinant proteins are able to bind single or double stranded DNA, an electromobility shift assay was used (EMSA). For this, 20 µl containing: 100 ng fluorescently labelled DNA or 200 ng non-labelled DNA (table S1), EMSA-buffer (10 mM Tris, 40 mM NaCl, 5 mM KCl, 5% glycerol, 2 mM MgCl_2_ and 0.025 mg/l BSA, pH 8), the recombinant protein (concentrations indicated in experiments) and water, was mixed and incubated at 20°C for 20 minutes according to the protocol of Chu et al. (45). EMSAS were performed using 6% native PAGE (1.5 kb fragment) or 10% (oligonucleotides). For detection, gels were stained with Midoori-Green for 20 minutes, and fluorescence signals were detected with a Typhoon scanner with 488 nm or 561 nm laser lines.

### Immobilization experiments

For testing of ComGC binding to ssDNA, biotinylated 12 nt polyA DNA was immobilized on streptavidin-coated agarose beads. Protein in 250 mM NaCl containing SEC-buffer (see above) was loaded onto DNA-bound beads. Washing steps 1-3 were performed by using SEC-buffer containing 250 mM NaCl. Elution was performed using SEC buffer containing 1 M NaCl. Samples were loaded with SDS-sample buffer.

To identify DNA binding proteins in competent cells lysate of a PY79 *Δrok* strain was used to increase the number of cells entering the competent state (K-state). As a negative control we used a strain which cannot enter the competence state and therefore not express the competence genes (*PY79, ΔcomK)* (table S3). For ssDNA experiments, a biotinylated primer was used, while for dsDNA fragments the same primer was complemented at 95°C for 5 minutes followed by a slow cooling (see list of oligonucleotides; table S6). For generating DNA-coupled agarose beads, streptavidin agarose resins were in a first step washed and equilibrated with annealing buffer (HEPES Buffer: 20 mM Hepes, 250 mM NaCl, KCl, 20 mM MgCl_2_ 0.01% tween) and subsequently BTN-DNA (biotin) was added to the beads to manufactures device and incubated for 1 h (IBA Live Science) at 4°C on a shaking platform. To generate *B. subtilis* lysate of competent cells, a culture in MC-media was grown to competence. Cells were harvested for 15 minutes and 4000 rpm at 4°C. Cells were resuspended in SEC-buffer (containing 0.06 µM Tween 20 and 0.1% Triton) to reach OD_600_20-25. Cells were disrupted by sonication for 20 seconds for a minimum of 5 times and in between each cycle cooled on ice. Cell debris was removed by centrifugation at full speed for 20 minutes. 400 µl of the supernatant was added to DNA coated beads and incubated for 1 h at 4°C on a shaking platform. Beads were washed three times with washing-buffer (HEPES Buffer: 20 mM Hepes, 250 mM NaCl, KCl, 20 mM MgCl_2_) and analysed *via* mass spectroscopy by the MarMass facility of Philipps University Marburg.

### Biolayer interferometry analysis (BLI)

For biolayer interferometry assays, a BLI-system (ForteBio, Pall Life Science) was used with High precision Streptavidin (ocet SAX2) biosensors (Sartorius, USA), which were derivatized with biotinylated DNA fragment (table S1). For BLI-measurements 20 mM Hepes, 250 mM NaCl, KCl, 20 mM MgCl_2_, 0.01% tween 20 (v/v) and 10 µM BSA was used. Biosensors were hydrated in buffer before use. A stable baseline was established first followed by association reactions with different concentrations of ComGD. Dissociation was measured by transferring the sensor to solely buffer. To remove immobilized protein from the biosensor, sensor was washed with buffer containing 20 mM Hepes, 1 M NaCl, 20 mM KCl, 20 mM MgCl_2._ Data was recorded by Blitz Pro 1.2.1.5 (ForteBio, Pall Life Science) software and analysed by Excel (Microsoft).

### Single-molecule tracking (SMT)

For single-molecule tracking, a derived protocol of Burghard-Schrod et al., 2022 was used (46). Cells were grown to competence in MC-media (see growth conditions). In order to facilitate uptake, cells were incubated for 1h at 37° and 200 rpm and further used for tracking. To prevent uptake of extracellular DNA cells were treated for 3 min at 37°C with 100 µg/ml DNAse I and washed. CY3-DNA or FAM-labelled-DNA was added to competent cells entering the stationary phase and incubated for 1 h in the dark (46). 3 µl of the culture was transferred to a round coverslip (25 mm, Marienfeld). 1% agarose pad (w/v). The pad was generated by sandwiching 100 μl of melted agarose between two 12 mm coverslips (Menzel, Germany), and subsequently, cells were pipetted onto the pad after removal or one coverslip. An inverted Nikon microscopy Ti Eclipse was used with an A=1.49 objective and an EMCCD camera (Hamatsu). Generated data were analysed by ImageJ (47). Analyses of bleaching curves revealed that the first 500 frames of the movies were to be removed, until 10% decline of the slope were reached. Oufti was used for cell meshes (48) and movies were analysed with Utrack after setting a minimal track length of 5 (49). The nature of tracks was then analysed by the SMTracker 2.0 software using SQD analyses, which describes the cumulative probability density function of squared displacements (50). The localization of errors was calculated by using the mean squared diffusion analyses, by taking the intercept of the graph with the y-axis.

### Sequence alignments and structural analysis

Sequence of respective proteins was sourced from Subtiwiki (https://subtiwiki.uni-goettingen.de/) or Uniprot (https://www.uniprot.or), and multiple sequence alignment tool MUSCLE https://www.ebi.ac.uk/) was used for the alignment of respective sequences of major/ minor pilins. Protein structures were produced by alpha fold 3 (https://alphafoldserver.com) and edited by Pymol (www.pymol.org). Formation of secondary structures was predicted by vector builder (DNA Secondary Structure Prediction Tool | VectorBuilder).

## Results

### The major pilin *Bs*ComGC forms monomers *in vitro*, while *Geobacillus* ComGC forms monomers and dimers

ComGC from *B. subtilis* has been shown to form higher oligomeric structures in protoplast supernatants (34). Therefore, it was hypothesized that this protein might constitute the core component of the extracellular *Bacillus* DNA uptake pilus, and was subsequently termed major pilin. Pilin proteins are notorious for being difficult to characterize *in vitro*, mainly because of to their partially lipophilic nature, due to the N-terminal transmembrane helix. Fig. 1 shows models for ComGC from *B. subtilis* (Fig.1A) and from *Geobacillus thermodenitrificans* (Fig. 1B). The 5 N-terminal amino acids are cleaved off by the pre-pilin peptidase ComC (51), the hydrophobic alpha helix remains within the lipid bilayer, and likely forms the core of the helical pilus assembly. In order to circumvent the problem of poor solubility yet to maintain the DNA-binding part, we constructed an N-terminal truncation of ComGC starting from amino acid 28 (red part Fig. 1A, red arrow Fig.1C) for heterologous expression in *E. coli,* using an N-terminal His-GST fusion to improve protein solubility.

**Figure 1:**
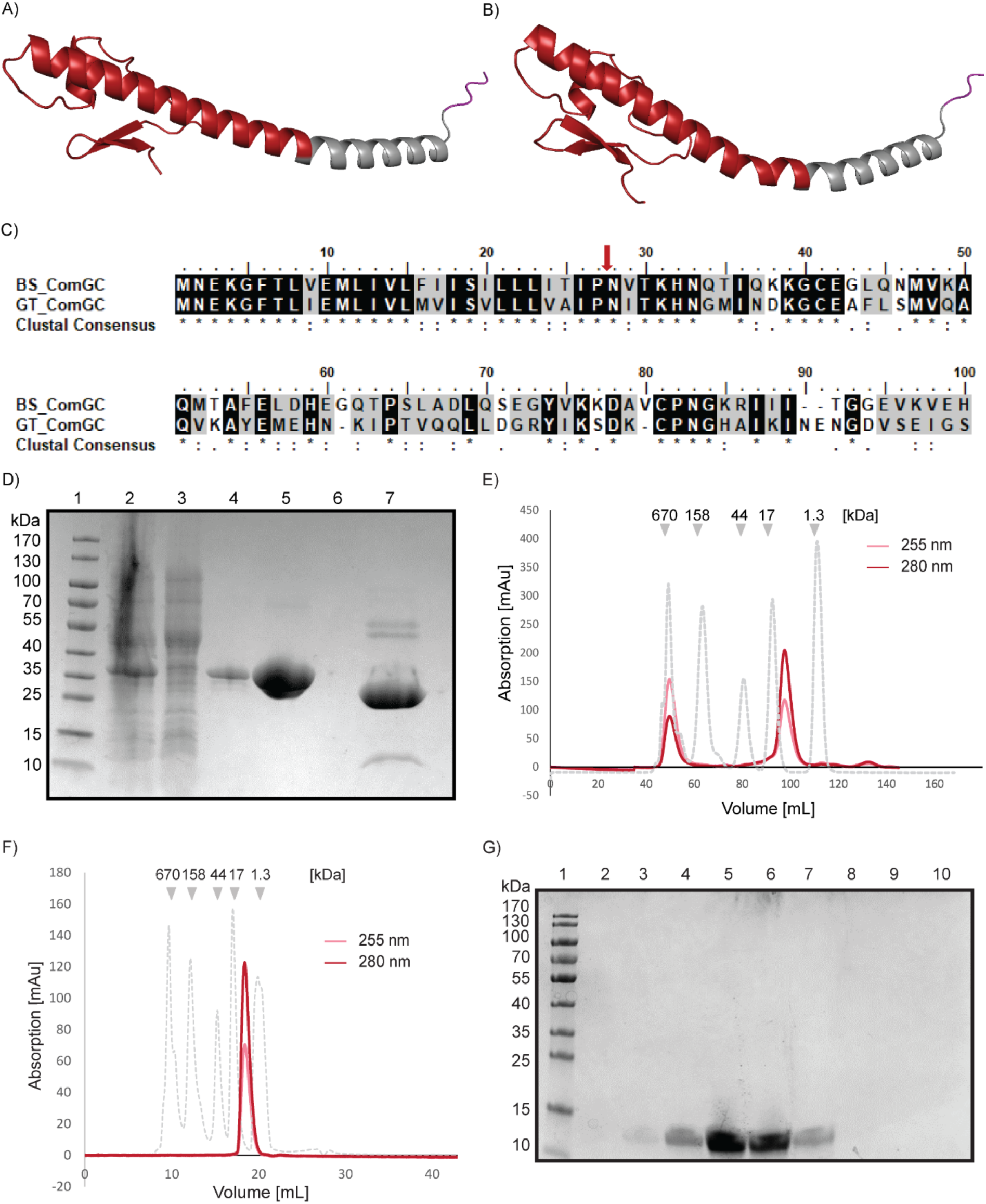
Structural and bioinformatic analysis of *Bacillus subtilis* and *Geobacillus thermodenitrificans* ComGC and purification and characterization of *Bs*ComGC. A) and B) show Alpha Fold structure-predictions for *Bs*ComGC (A) and *Gt*ComGD (B). Red colour indicates the hydrophilic part of the protein used for purification. Grey colour indicates part of the transmembrane part and purple the N-terminal region that is processed by the pre-pilin peptidase ComC. C) Multiple sequence alignment of *Bs*ComGC and its homologue *Gt*ComGC. Identical amino acids are indicated in black, similar in grey and white displaying no homology. Alignment was done using MUSCLE (52) and represented by Bioedit. Red arrow indicates starting amino acid sequence used for purification. D) Coomassie stained SDS PAGE of Ni-NTA purification steps of His-GST-*Bs*ComGC. 1) Prestained pageruler (Thermofisher scientific), 2) lysate, 3) flow through, 4) wash, 5) elution and 7) elution treated with TEV-protease. E) Chromatogram of size exclusion chromatography of elution fractions of ComGC. F) Chromatogram of analytical size exclusion chromatography of monomeric fraction of *Bs*ComGC. Red colour absorption at 280 nm, pink colour absorption at 255 nm. Standards are shown in dashed grey line. G) Coomassie stained SDS PAGE of monomeric fractions after size-exclusion. Lane 1 represents pre-stained page ruler, lane 2-7 monomeric fractions of ComGC.

His-GST-*Bs*ComGC was purified by Ni-NTA-affinity chromatography (Fig. 1D). The eluted protein migrated at a size of 37 kDa corresponding to the size of full-length His-GST-ComGC (Fig. 1D). The affinity/solubility tag includes an ENLYFQS TEV-recognition site distal to the tag and proximal to the protein of interest, so that TEV-protease can remove the solubility tag. Elution fractions were treated with TEV-protease followed by reverse Ni-NTA chromatography to separate the His-GST-tag and the TEV protease from ComGC (Fig.1D). Size exclusion chromatography (to remove impurities and aggregated proteins) showed two peaks: proteins eluting close to the void volume, corresponding to 670 kDa, and a second fraction at around 100 ml corresponding to monomeric ComGC (∼10 kDa) (Fig. 1E and 1G). To gain more insight into the oligomerization of *Bs*ComGC we used analytical size exclusion chromatography, in which the purified monomeric fraction of ComGC were re-run. Again, *Bs*ComGC eluted at a size corresponding to monomeric ComGC (Fig. 1F), showing that this is the major form of truncated ComGC *in vitro*.

In order to assess differences between *Bs*ComGC with homologs in other *Bacilli* we constructed strains expressing *Geobacillus thermodenitrificans* His-MBP-ComGC (hereafter named *Gt*ComGC), which is closely related to *Bs*ComGC and shows high sequence - (Fig. 1C) and structure conservation (Fig. 1A and B). We also used a similar truncation for heterologous expression of *Gt*GomGC in *E. coli* cells (Fig. 1B, red part). In case of *Gt*ComGC a His-MBP-tag fused to the protein resulted in sufficient solubility of the protein (50 kDa, Fig. 2A).

**Figure 2:**
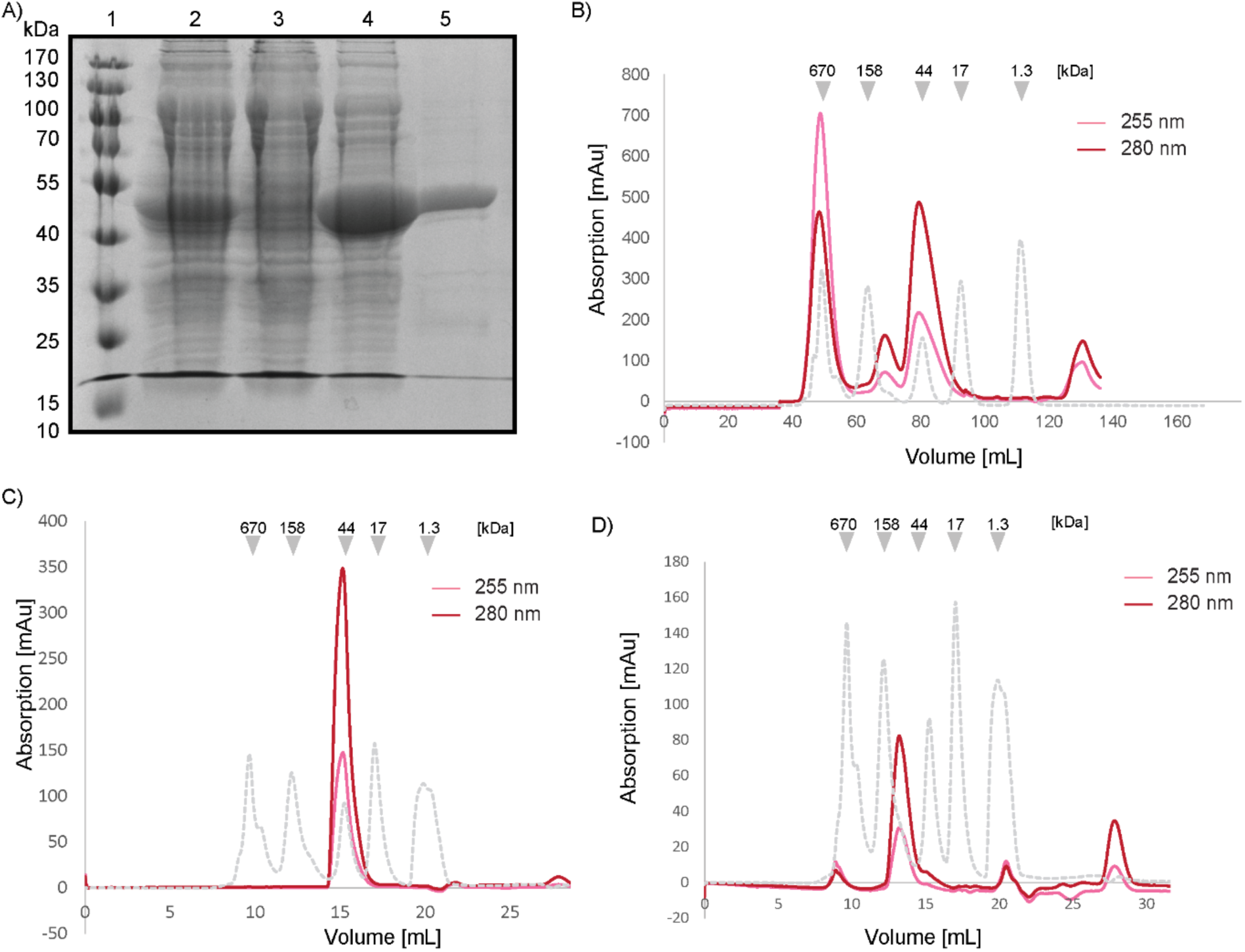
Purification and characterization of *Gt*ComGC. A) Coomassie stained SDS-PAGE of Ni-NTA purification steps of His-MBP-*Gt*ComGC. 1) Prestained page ruler, 2) lysate 3) flow through 4) wash and 5) elution. B) Chromatogram of size exclusion chromatography of elution fraction of His-MBP-*Gt*ComGC. C) Chromatogram of analytical size exclusion chromatography of monomeric fraction of His-MBP-*Gt*ComGC. and D) chromatogram of dimeric elution fraction of His-MBP-*Gt*ComGC. Red colour absorption at 280 nm, pink absorption at 255 nm (B, C and D). Standards are shown in dashed grey line.

In contrast to His-GST-fusions of *Bs*ComGC, His-MBP was not cleaved due to the low solubility of *Gt*ComGC (data not shown). Size exclusion chromatography was efficient in removing the impurities (Fig. 2B). Chromatograms showed 3 defined peaks above 1.3 kDa, all of which contained His-MBP-*Gt*ComGC (Fig. 2B, Fig. S1B). Peaks correspond to monomeric (50 kDa) and dimeric (100 kDa) His-MBP-*Gt*ComGC, which was also present in the void volume, possibly forming multimers (Fig. S1B). Please note that hereafter, the His-MBP tag will not be explicitly stated for *Gt*ComGC.

To further investigate oligomerization of *Gt*ComGC, fractions were analysed by analytical size exclusion. A sample generated from the proteins isolated from the third peak of *Gt*ComGC (monomer) eluted at 16 ml corresponding to monomeric *Gt*ComGC (Fig. 2C), proteins retrieved from the second peak (dimeric) *Gt*ComGC, largely eluted within the dimer peak (Fig. 2D). These analyses show that *Gt*ComGC forms stable monomers and dimers. The fact that the *Gt*ComGC monomers remained monomeric shows that the His-MBP tag did not cause dimerization or multimerization artefacts.

### The minor pilin *Gt*ComGD is present in an equilibrium between monomeric and dimeric states

Besides the major pilin ComGC, the competence pili generally contain minor pilins; in case of *B. subtilis,* these are ComGD, ComGE, ComGG, and based on bioinformatical analysis, likely also ComGF (53). The presence of multiple, closely related pilins in a single structure could be explained by either functional redundancy or specialized functions of the homologs. Several studies have indicated that minor pilins form a complex that initiates T4P assembly (54–56). If such initiation mechanism was conserved for all type IV filaments, the presence of all individual minor pilins would be generally necessary for pilus formation. In addition to their role in pilus formation initiation, minor pilins might contribute specialized attributes to the pilus structure by modifying polymer stability and binding affinity. We chose to characterize one member of the minor pilins in order to compare its properties with those of major pilin ComGC.

*G. thermodenitrificans* possesses at least two minor pilin genes, encoding for proteins showing homology to ComGD and ComGF (57). According to the KEGG database (https://www.kegg.jp), the downstream genes are also annotated as homologues of minor pilins, and despite low homology could correspond to minor pilins *Bs*ComGE or *Bs*ComGG.

Sequence alignment and structure analysis of *Bs*- and *Gt*ComGD are shown in Fig 3A-C. We were able to purify *Gt*ComGD fused to the His-MBP-tag to high concentrations (Fig. 3D). Due to the limited solubility of the cleaved product, we performed experiments with the fusion protein. Preparative size exclusion showed 3 defined peaks (Fig. 3E), corresponding to void volume, dimeric and monomeric His-MBP-*Gt*ComGD (at 50 kDa, dimers at 100 kDa) (Fig. S1C). This is indicative of the formation of an equilibrium of mono-as well as dimers of His-MBP-*Gt*ComGD. When run on analytical gel filtration, both monomeric and dimeric fractions of His-MBP-*Gt*ComGD split up into 2 peaks (monomers and dimers), characteristic of a dynamic equilibrium (Fig. 3F and G). Hereafter His-MBP-*Gt*ComGD is termed *Gt*ComGD.

**Figure 3:**
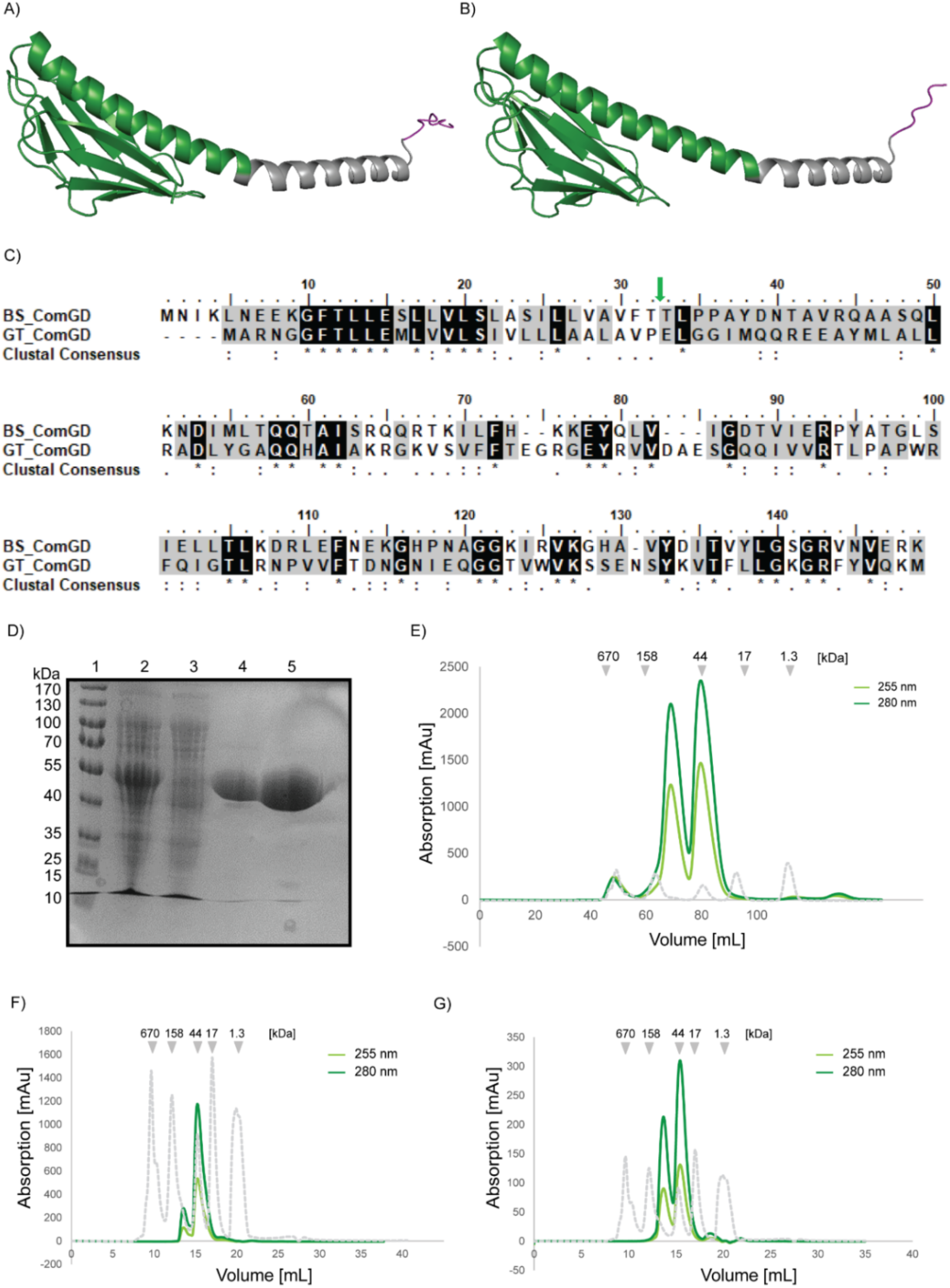
Structural and bioinformatic analysis of *Gt*ComGD and purification and characterization of His-MBP-*Gt*ComGD. A) and B) show alpha fold models of *Bs* (A) and (B) *Gt*ComGD structures. Green colour indicates soluble, purified part of the protein, grey colour indicates part of the membrane part and purple the N-terminal region that is cleaved off. C) Multiple sequence alignment of *Bs*ComGD and its homologue *Gt*ComGD. Identical amino acids are in black, similar ones in grey; green arrow indicates the start of the purified protein. Alignment was done using MUSCLE (52) and represented by Bioedit. D) Coomassie stained SDS-PAGE of Ni-NTA purification steps of His-MBP-*Gt*ComGD. Lanes represent 1) Prestained page ruler, 2) lysate, 3) flow through, 4) wash and 5) elution fraction. E) Chromatogram of size exclusion chromatography of Ni-NTA elution fractions of His-MBP-*Gt*ComGD. F) Chromatogram of analytical size exclusion chromatography of monomeric fraction of His-MBP-*Gt*ComGD and G) dimeric fraction. Light green absorption at 255 nm, dark green at 280 nm. Size standards are indicated by dashed grey line.

### Minor pilin *Gt*ComGD and major pilin ComGC are able to bind ds-as well as ssDNA *in vitro*

Competence pili of *Streptococcus* were shown to bind dsDNA *in vivo*, along their entire length, but it is unclear by which subunits of the pilus (42). In *S. sanguinis,* major pilin ComGC does not bind to DNA, but loss of minor pilins completely abolishes DNA uptake *in vivo* (58). To test if *Bacillus* major pilins could be directly involved in DNA binding, we tested DNA the binding capability of purified His-MBP-*Gt*ComGC and *Bs*ComGC using electromobility shift assays (EMSA), employing DNA fragments of various lengths. A dsDNA fragment of 2400 bp was bound by *Bs*ComGC within the low µM range (Fig. 4A), and likewise a 50 bp dsDNA fragment. Efficient binding of the large fragment occurred rather sharply between 60 and 150 µM (Fig. 4A), indicating cooperative binding to DNA. Likewise, His-MBP-*Gt*ComGC showed clearly shifted DNA bands (Fig. S2A). DNA constructs were shifted irrespective of their sequence, indicating unspecific binding of DNA. Experiments were done in the presence of at least 300 mM salt, ruling out low-affinity binding.

**Figure 4:**
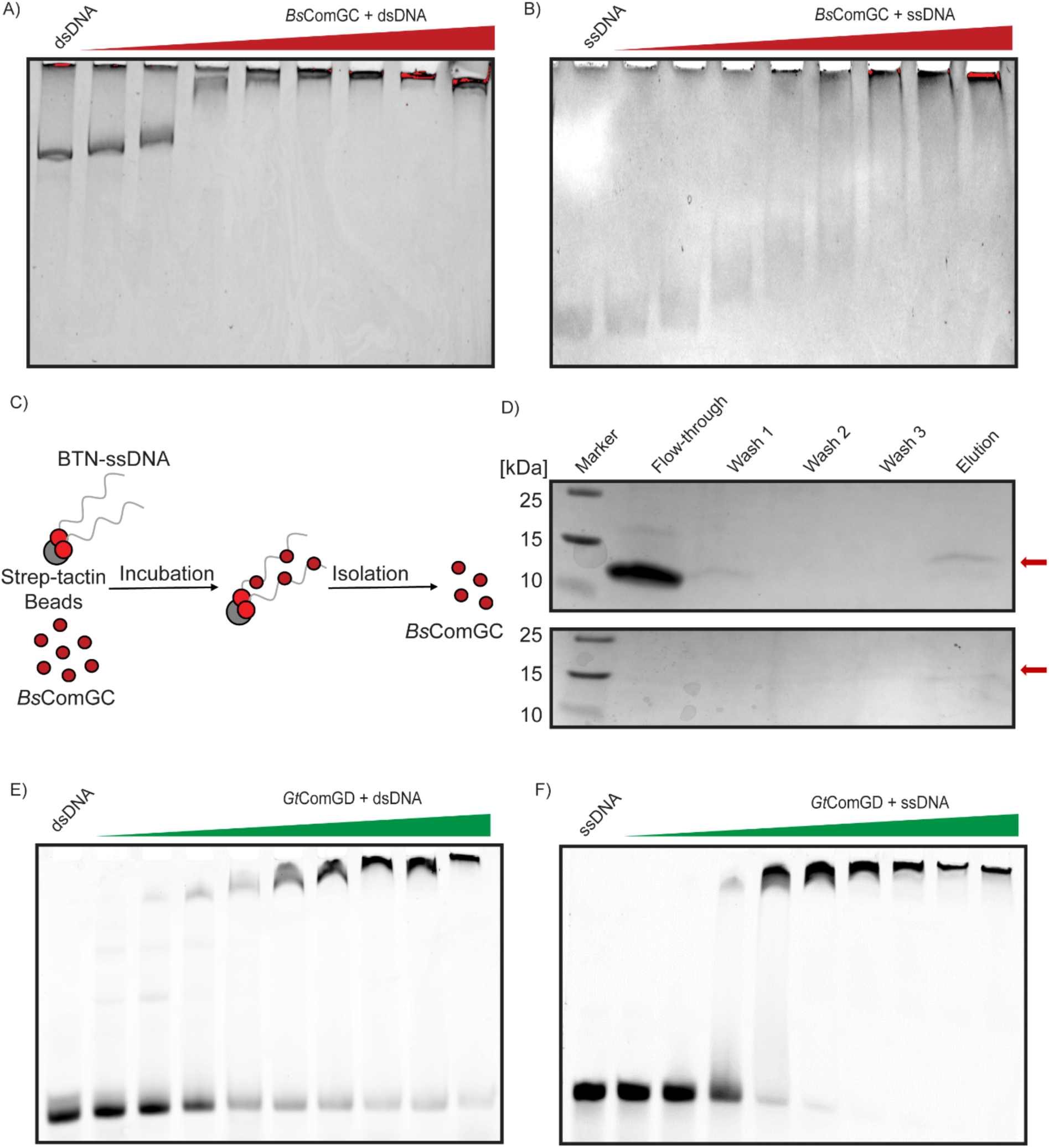
DNA binding of major and minor pilins to ds- and ssDNA. A) Midoori-stained native PAGE (electromobility shift assays) of *Bs*ComGC and 2400 bp dsDNA (B) or 40 bp ssDNA (Fig. S3A). Protein concentrations of *Bs*ComGC: 0 (lane 1), 30/ 60/ 150/ 320/ 450/ 600/ 850/ 900 µM (lanes 2 to 9), triangle indicates increasing protein concentrations. C) Schematic representation of the experimental setup of figure D. Biotinylated ssDNA coated streptactin beads were incubated with purified protein *Bs*ComGC. D) SDS-PAGE of *Bs*ComGC binding to immobilized 12 nt polyA ssDNA, with a high concentration of *Bs*ComGC (370 µM, upper panel) or low concentration (120 µM lower panel). E) Native PAGE of electromobility shift assays of E) FAM labelled 40 bp dsDNA and F) Native PAGE of FAM-labelled 40 base ssDNA (Fig. S3A) of *Gt*ComGD. Protein concentrations of *Gt*ComGD ranged from 0 (lane 1) to 1.8/ 4/ 10/ 18.5/ 25/ 37/ 48/ 55/ 60 µM (lanes 2 to 10). Gels were detected by a fluorescent imager.

In order to test if pilins may also bind to ssDNA, we used a 40 base oligonucleotide that was predicted to lack any double strand regions (Fig. S3A). Interestingly, major pilins also bound to the ssDNA fragment (Fig. 4B). Here, *Bs*ComGC/ssDNA complexes increased steadily in size, suggesting non-cooperative binding (Fig 4B), while *Gt*ComGC formed distinct high molecular weight complexes (Fig. S2A). Due to the nature of the shifts, a true KD value is difficult to calculate; for *Bs*ComGC, is lies around 100 µM based on a clearly visible change in binding efficiency between 60 and 150 µM shown in Fig. 4B. Thus, binding to double – as well as single-stranded DNA occurred within a similar range of protein concentration. Because ssDNA can be strongly bent, ssDNA binding activity of major pilins could allow for efficient uptake of ssDNA or of DNA containing ssDNA regions.

To further test the ability of *Bs*ComGC to bind single stranded DNA we used an affinity immobilization experiment assay. For this, 12 nt biotinylated polyA-ssDNA (Fig. S3B) was bound to Streptavidin-coated agarose beads, and was incubated with purified *Bs*ComGC. After washing steps using 250 mM NaCl, proteins were eluted using 1 M NaCl. While the protein alone was unable to bind to non-modified beads (Fig. S2G), *Bs*ComGC could be detected in the elution fraction (Fig. 4D, lower gel) and also in the washing steps when a high protein concentration was used (Fig. 4D upper gel). These data further prove the capability of ComGC to bind to ssDNA.

Minor pilins of *Legionella pneumophila* have been reported to bind dsDNA *in vitro* (59). To test the ability of the minor pilin ComGD to bind DNA we performed EMSA with purified His-MBP-*Gt*ComGD. Although for *B. subtilis* no specific DNA-uptake-sequences have been reported, we used a variety of endo-as well as exogenous DNA sequences to assess specific or non-specific DNA binding. As for the major pilins, we observed increasing formation of slow-migrating complexes with increasing concentrations of the protein (Fig. 4E), implying the ability of ComGD binding dsDNA *in vitro*. The nature of the shifts implies lack of cooperativity in binding to dsDNA, and a K_D_ value in the range of 20 µM (between lanes 5 and 6, Fig. 4E).

Because of the surprising ssDNA-binding activity of major pilins, we assessed the ability of the minor pilin (ComGD) to bind to ssDNA. Interestingly, His-MBP-ComGD shifted ssDNA (40 nt, Fig. S3A) with even higher apparent affinity than dsDNA (Fig. 4F), based on more efficient binding at lower protein concentrations. Again, binding was sequence independent. A strong change in DNA shifting occurring between 10 and 18.5 µM (lane 4 and 5, Fig. 4F) of *Gt*ComGD suggests cooperative binding to ssDNA, and a KD value in this µM range. We employed a more quantitative technique for calculating K_D_ values for ComGD (see below). DNA binding of purified monomeric or dimeric fractions of ComGD was indistinguishable (Fig. S2 B,C), showing that both forms of ComGD are equally able to bind to ssDNA.

### ComGD binds to ssDNA and dsDNA with similar affinity

In order to quantitatively analyse the DNA binding capability of pilins we used biolayer-interferometry (BLI)-analysis with biotinylated single - or double stranded DNA attached to the sensor (schematic illustration in Fig. S4A). To exclude unspecific binding of the protein to the sensor, we measured the wavelength shift when solely adding His-MBP-ComGD to a biosensor (Fig. S4C). No wavelength shift was detected, in contrast to biotinylated DNA, which showed a shift when binding to the strep-tag-modified sensor, as expected (Fig. S4C). To additionally rule out unspecific binding of the His-MBP-tag itself to the biosensor, we tested the His-MBP-tag for DNA binding activity, and did not detect any binding to DNA-coated sensors (Fig. S4B).

When ComGD was added to DNA-loaded sensors using 2000 bp dsDNA, *Gt*ComGD showed a concentration-dependent increase in wavelength shift (Fig. 5A). The protein dissociated after a washing step with salt (250 mM). ComGD also bound to a 48 nt hairpin DNA (Fig. 5B), predicted to form mixed single – and double stranded regions (Fig. S3D). By plotting the dependency of shifts relative to the concentration of the protein, association constants were determined (Fig. 5A and B, right panel). For hairpin DNA we determined a K_D_ of 98 µM (Fig. 5), while for 2000 bp dsDNA we determined a constant of 96 µM (Fig. 5B), which do not differ significantly. Therefore, length of dsDNA does not appear to influence binding of purified minor pilin.

**Figure 5:**
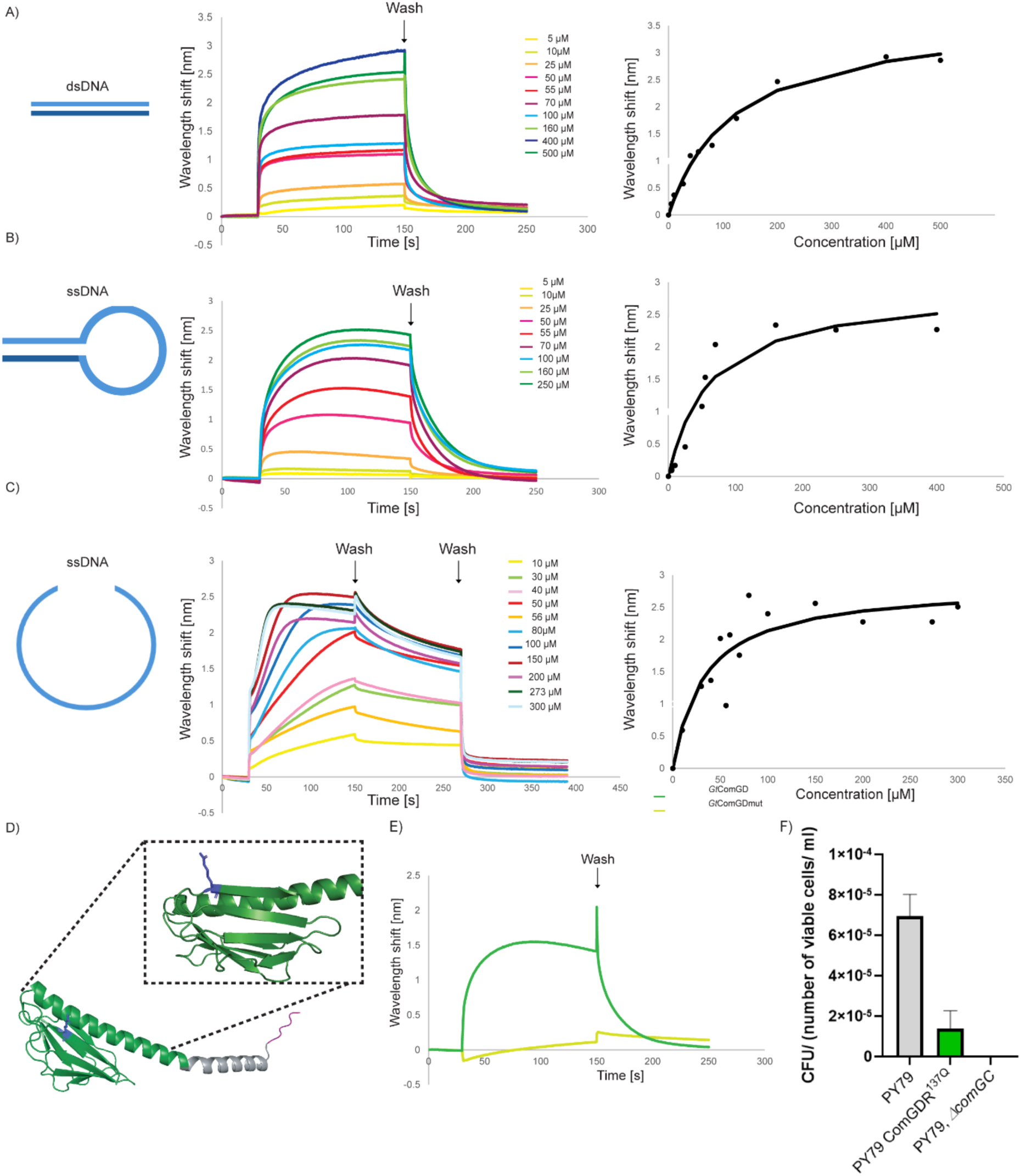
Biolayer interferometr*y* analysis of the interaction of ds - and ssDNA with *Gt*ComGD. A-C) BLI sensors loaded with biotinylated ss-and dsDNA were incubated with increasing concentrations of *Gt*ComGD (shown in the inset). Left panels indicate the structure of DNA on the sensor. Middle panel graphic views of kinetic interactions, resulting from wavelength shifts during the association and dissociation phases. Right panels are representative graphs of wavelength shift dependent on added protein concentrations. Arrow indicates washing step with buffer (250 mM NaCl) or the second wash step with 1 M NaCl). D) Alpha fold model of *Gt*ComGDR139Q. The mutated arginine (R) is indicated in blue. Dashed lines indicate zoomed region. E) Representative measurement of association and dissociation kinetics of His-MBP-*Gt*ComGD wild-type protein (dark green) or His-MBP-*Gt*ComGD variant (light green). F) Transformation efficiency assays (CFUs/number of viable cells per ml) using a strain expressing mutated ComGD (R139Q). Error bars indicate SD. Graph generated by GraphPadPrism.

To confirm and quantify ssDNA binding we tested an oligonucleotide predicted not to form any secondary structures (Fig. S3A). Also, with this DNA fragment we were able to detect a concentration-dependent wavelength shift, and interestingly, we detected very slow dissociation of ComGD from the DNA during the washing step with low salt concentration. Instant dissociation occurred at 1 M NaCl (Fig. 5C). This shows that *Gt*ComGD binds to single stranded DNA with higher affinity than dsDNA, reflected in a lower K_D_ value of 53 µM (Fig. 5C). These data corroborate with EMSA experiments that suggested higher affinity to ssDNA than to dsDNA (Fig. 4E and F).

In order to obtain insight into the nature of ssDNA binding by pilins, we sought to identify residues involved in binding. We mutated several positive residues in ComGD that are conserved in minor pilins, and tested mutant versions for DNA binding ability. Protein variants were purified under the same conditions as wild-type protein (Fig. S5A-F). Size exclusion chromatography revealed formation of monomers and dimers like wild-type His-MBP*-Gt*ComGD (Fig. 3), indicative of proper folding of protein variants. For testing ssDNA binding ability, BLI-analysis was performed. By exchanging a conserved Arginine (R) at position 139 (note *Gt*ComGD is 5 amino acids shorter than *Bs*GomGD, Fig. 3C) to Glutamine (Q) (Fig. 5D), neither single nor dsDNA binding activity could be measured (Fig. 5E), at the same concentration where wild-type protein exhibited strong DNA binding (dark green curve, 100 µM). Mutation of a Lysine (K123) on the beta-sheet did not prevent ss- or dsDNA binding compared to the wild-type. From this analysis, we can assume that R139 is involved in DNA binding, supporting the notion that pilins have a genuine ssDNA binding surface.

To test if the mutation has an effect of DNA uptake *in vivo* we used CRISPR-cas9 to generate a strain expressing the corresponding protein variant *in B. subtilis.* By performing transformation efficiency assays, we obtained transformation efficiencies of 1.4 (+/- 0.9) x 10^-05^ CFU/ml, compared to 6.8 x (+/- 1.2) x 10^-05^ CFU/ml for wild type cells (Fig. 5F), and thus a roughly 5-fold reduced transformation rate. Thus, the lack of R139 that is highly important for DNA binding in ComGD translates into to a considerable defect in transformability *in vivo*. This is in accordance with previously published studies for which a deletion of the homologous residue in *Streptococcus sanguinis* as well as *Streptococcus pneumoniae* also leads to reduced DNA uptake *in vivo* (54,55).

We also mutated several positive residues in ComGC (see Fig. S6 for purification of soluble variants) and tested them for DNA binding ability by EMSAs but could not identify any single mutation that prevented DNA binding (Fig. S2E, F) in comparison to the wild-type BsComGC (Fig. S2D).

### Major pilin ComGC can be enriched from *B. subtilis* cell extracts using ssDNA

In order to move our analyses towards *in vivo* experiments, we reasoned that if pilins were indeed able to bind to ssDNA in cells grown to competence we would expect that they could be co-purified with ssDNA added to a culture of cells during the K-state. To this end we incubated beads coated with 40 base ssDNA (Fig. S3G) and added extract from *B. subtilis* cells grown to competence for 60 minutes, using cells carrying a *rok* deletion to increase the percentage of cells that are entering the K-state, or *comK* mutant cells as control. After washing of the beads, bound molecules were identified using mass spectrometry (Fig. 6A). The results of proteins of interest are shown in table S1, a volcano plot of identified proteins is shown in Figure S7. As expected from their known ability to bind to ssDNA with high avidity, unique peptides of SsbA (acts at replication forks) and SsbB (competence-specific SSB) were identified at high frequency, of RecA, a multifunctional protein involved in homologous recombination and DNA repair, as well as of DprA, a protein that hands over incoming ssDNA to RecA (60,61). Periplasmatic receptor protein ComEA, ComGC and minor pilin ComGF were also found as significantly enriched (Fig. S7). In contrast, SsbB, ComGC, ComGF and ComEA (but not SsbA) were absent from eluates of *comK* mutant cells (Fig. S7 and table S1).

**Figure 6:**
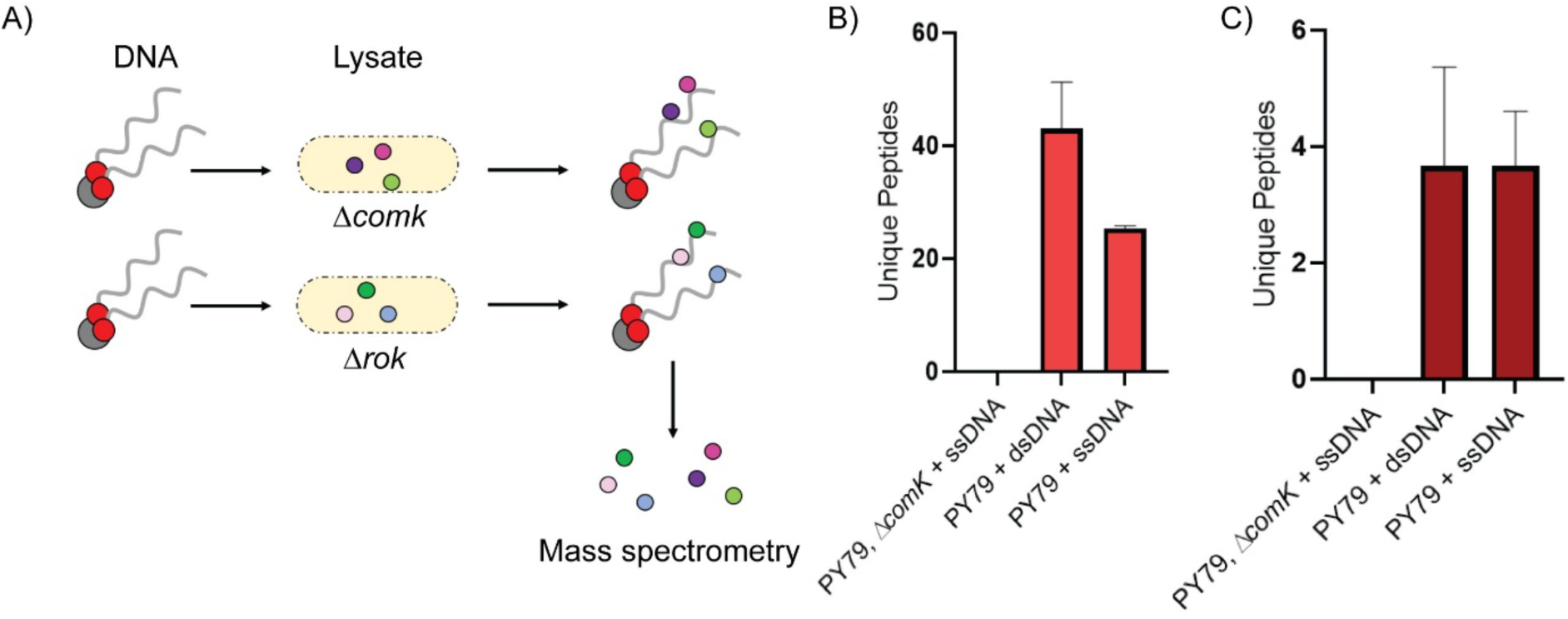
Immobilization experiments by using single or double stranded DNA as a bait. A) Schematic illustration of experimental set up. Biotinylated single or double stranded DNA was immobilized on Strep-tactin beads and incubated with lysates of a *comK* or a *rok* mutant strain. Proteins that were bound to BTN-ss/dsDNA coated beads were analysed by mass spectrometry. B) and C) showing average number of unique peptides from 6 experiments of ComEA (B) or ComGC (C), compared to average number of peptides from 3 experiments using *comK* mutant cells (table S1).

In order to obtain more quantitative data, we analysed the number of unique peptides of ComEA and ComGC that were enriched using beads coated with ss- or dsDNA (listed in table S1). (Fig. 6). ComEA has been characterized to feature higher affinity to double stranded DNA than to ssDNA (62). Indeed, this was reflected in our experiments (Fig. 6B). Additionally, we detected nuclease YhaM that was recently reported to be involved in DNA uptake *in vivo*. YhaM shows high affinity to double-as well as single stranded DNA *in vitro,* and even higher affinity to ssDNA (63), which was also seen in our experiments by elevated unique peptide counts (table S1). Contrarily, unique peptides of ComGC showed similar peptide counts for single as well as double stranded DNA used as a bait (Fig. 6C), supporting the idea of ss- and dsDNA-binding affinity within a similar range. Of note, minor pilins other than ComGF may not have been detected based on a putative low copy number within competence pili.

### SsDNA accumulates within the *B. subtilis* periplasm in a manner analogous to dsDNA

Based on our *in vitro* experiments, we would predict that *Bacillus subtilis* cells grown to competence should take up double - as well as single stranded DNA into the periplasm. It is well characterized that *Bacillus subtilis* moves double-stranded DNA from the environment through the cell wall into the periplasms where it accumulates (46,64), converts taken-up DNA into ssDNA and translocates it into the cytosol for homology-driven recombination (4).

In order to study the motion and dynamics of ssDNA *in vivo*, we used single molecule tracking with CY3-ssDNA fragments forming secondary structures (“48 bp ssdsDNA” Fig. 7A, Fig. S3E), as well as single stranded DNA labelled with FAM solely forming single stranded regions including various lengths of DNA (Fig. S3A, S3C). In addition to visualizing the preferred localization of molecules with a precision of well below 50 nm, SMT allows to obtain information on the dynamics of molecules and thus existence of possibly different states of mobility, which gives insight into formation of molecular complexes in live cells.

**Figure 7:**
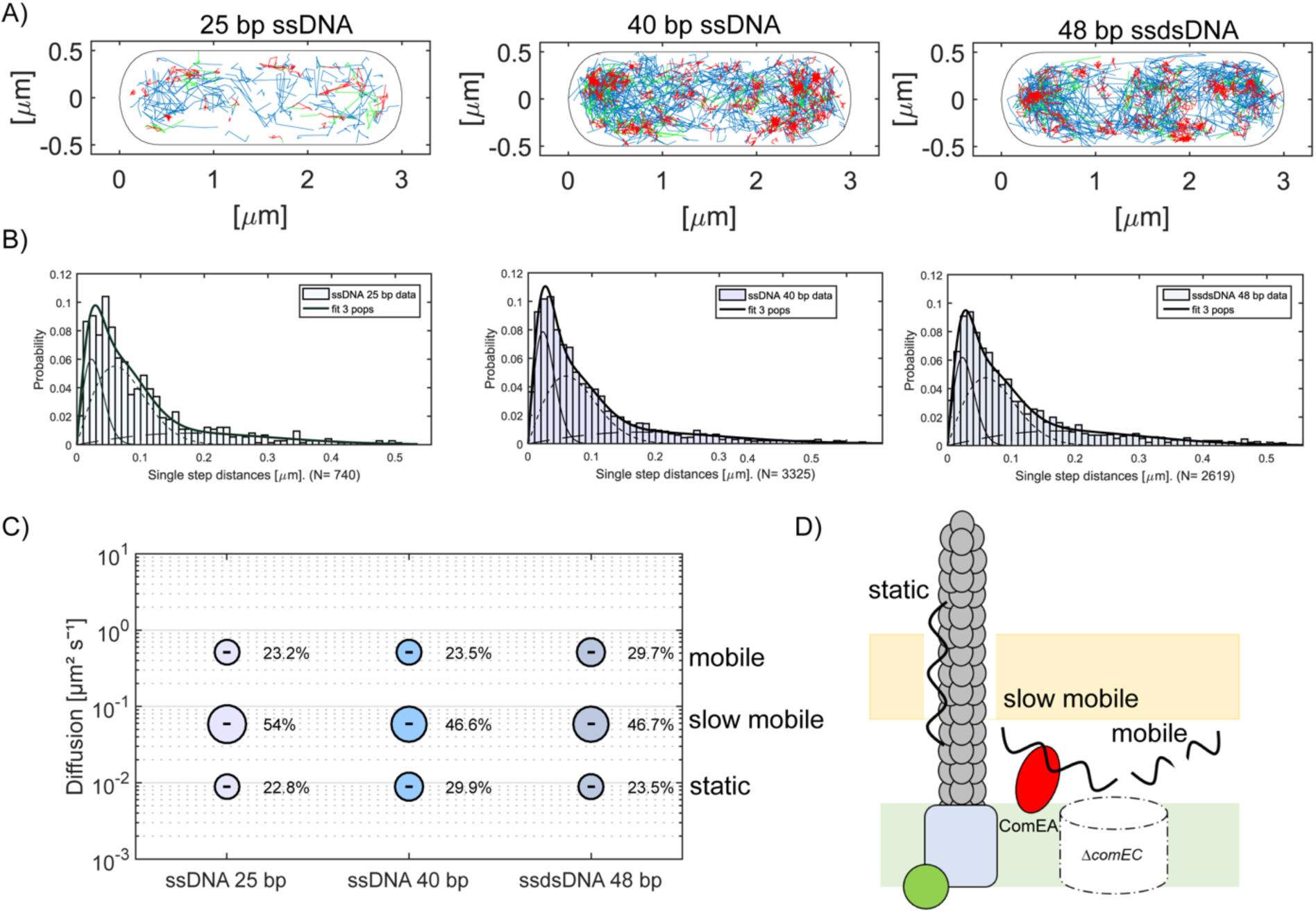
Single molecule tracking of PY79, *ΔcomEC, Δrok* cells incubated with ssDNA. A) Confinement maps showing the tracks on a standardized cell of ***Δ****comEC* ***Δ****rok* cells incubated with CY3- or FAM-labelled single stranded DNA. Lengths of the used DNA fragments are indicated above the panels. Red = confined tracks, blue = freely diffusive tracks, green = tracks showing transitions between confined and free diffusion. 25 bp DNA is indicated in light blue, 40 bp in blue and 48 bp in grey/blue. B) Squared displacement (SQD) analyses; histograms show the probability of the single step distances of acquired data. Solid thin line: static population, dotted line: medium-mobile population, dashed line: fast-mobile fraction; thick solid line combined three population fit. C) Bubble plots of acquired data from three independent biological replicates showing the fraction sizes of labelled ssDNA and their average diffusion constants in competent *B. subtilis* cells (table S3). D) Cartoon of the fractions of fluorescently labelled DNA. T4P is indicated in grey, ComEA (red), ComGA (green), ComEC in white (dashed line indicates deletion of *comEC*), ComGB (light blue).

We incubated *B. subtilis* cells grown to competence with the labelled ssDNA according to the protocol from Schrod et al., extended the incubation time from 30 minutes to 1h (46), followed by DNase treatment for 3 minutes at 37°C. By using a Δ*rok* strain with an additional *comEC* deletion, taken up DNA would remain in the periplasm for easier tracking. Fig. 7A shows that indeed, observed tracks were mostly observed at the periphery of cells; ssDNA of a length of 25 bases was taken up only very inefficiently (table S2, 1.5 tracks per cell compared with 3.8 for 40 nt ssDNA). Confined tracks of ssDNA represent molecules remaining within an area of 106 nm (2.5-fold the localization error), i.e. molecules that are not freely diffusing. ssDNA was found in this state at many positions within the cell’s periphery, same as freely diffusing molecules (blue) and molecules changing between both types of motion indicated in green.

Squared displacement analysis (SQD) describes the cumulative probability function molecules found in certain diffusive states. The data are depicted as jump distances in Fig. 7B, showing the probability of molecules moving with certain lengths of steps. Three Rayleigh fits were required to adequately describe the data, which was previously also described for dsDNA (46). The DNA molecules of 40 nt ssDNA (forming no secondary structures) had diffusion constants of 0.009 µm^2^/s for the static fraction, 0.06 µm^2^/s for the slow-mobile fraction and 0.51 µm^2^/s for the fast-mobile fraction. The latter constant is characteristic of a (large) freely diffusing (globular) molecule, and although the ssDNA molecule has only a weight of about 20 kDa, a large polymer will have a lower diffusion constant than a globular protein of the same size. About 23.5% of molecules were in this state (Fig. 7C). 46.6% of molecules showed low mobility, likely ssDNA bound to ComEA, which also shows such low mobility within the periplasm (65). The static population of 29.9% (table S3) of ssDNA molecules may consist of ssDNA molecules bound to pili during binding/retraction (schematic illustration 7D). These data are comparable to previous experiments tracking labelled dsDNA (46). Tested DNA fragments of 25 bp ssDNA and 48 bp ssDNA forming secondary structures revealed comparable sizes of static, slow-mobile or mobile fractions (Fig. 7C). In order to better compare molecule mobilities, fits having similar diffusion constants were used for SQD analyses. All tested DNA fragments showed a comparable behaviour (Fig. 7, table S4 shows small differences between molecules using individual fitting), independent of the formation of hairpin structures, or the length or the sequence of the DNA. Thus, ssDNA is efficiently taken up into the periplasm and moves within this compartment in a manner highly comparable to dsDNA.

## Discussion

Over previous decades, multi-antibiotic-resistant strains of bacteria have spread, which is a major problem for human health care systems worldwide. The reason for the rapid spread of genes empowering antibiotic resistance is efficient horizontal gene transfer between bacteria, which plays a major role in bacterial evolution (4). A key mechanism of HGT is natural competence, through which bacteria actively take up environmental DNA, e.g. antibiotic-resistant genes, and incorporate them into their chromosomes. Starting with the discovery of isolated DNA being able to convert a non-pathogenic bacterium into an infective species (66), work on transformation via competence has focussed on double stranded DNA being the transforming agent, and uptake of ssDNA was rarely taken into consideration. However, uptake of or transformation via ssDNA has been detected for different competent bacterial species (67,68). We have tested this general idea by analysing DNA binding properties of competence pili, the machinery initially interacting with environmental DNA, from the soil bacterium *Bacillus subtilis*, which is closely related to pathogenic species such as *B. anthracis*, and serves as a model organism for natural competence. We show that *B. subtilis* major pilin and *Geobacillus thermodenitrificans* major and minor pilins are able to bind to double as well as to single stranded DNA, and that ssDNA is efficiently taken up into the periplasm. Our data suggest that import of ssDNA, be it as food source or transforming agent, has been underestimated.

In all bacteria studied in detail so far, uptake of environmental DNA is dependent on a cell envelope-spanning machinery composed of type IV pili that capture and guide DNA through the multi-layered peptidoglycan in Gram-positive species or outer membrane and cell wall in Gram-negative bacteria, by recurring extension and retraction cycles. How these “fishing rods” bind to environmental DNA in detail is a matter of current research. Pili contain major and minor pilins, the latter are thought to form a pilus tip-complex that binds to DNA (19,41,59). It is not clear yet if minor pilins are only present at the pilus tip, or also along the length of the pilus, mixed with the major pilin. Previous studies showed for *S. pneumoniae* that dsDNA is bound along the entire length of pili (42), while DNA binding in *Streptococcus* exclusively occurs via minor pilins, which do not show DNA binding activity when purified (54,55). On the other hand, minor pilins FimT from *Legionella pneumophila* or from other Gammaproteobacteria show DNA binding *in vitro* (59). Our *in vitro* experiments clearly demonstrate that in *B. subtilis* and *G. thermodenitrificans* major pilins ComGC as well as *G. thermodenitrificans* minor pilin ComGD are able to bind dsDNA *in vitro*, showing that major as well as minor pilins can possess DNA binding capacity.

The cell wall is a tight meshwork of crosslinked peptidoglycan strands that must withstand internal osmotic pressure of 2 (in Gram-negatives) to more than 10 (in Gram-positives) atmospheres, wherefore the Gram-positive cell wall has a multitude of layers. It follows that the wall cannot contain larger discontinuities, such that pili - having a diameter of about 8 nm - will not have more space to pull DNA through the wall upon retraction. The detailed mechanisms of this transport has been an open question, particularly with respect to the spatial restrains in channeling double stranded DNA with a persistence length of hundreds of base pairs through a narrow pore. A key finding of this work is that the major pilin of *B. subtilis* and its homologue from *G. thermodenitrificans* as well as its minor pilin, bind to partially single stranded DNA *in vitro* with a similar affinity (of ∼ 98 µM) than to dsDNA (∼96 µM). SsDNA forming no secondary structures bound in the same order of magnitude, with a lower binding constant (53 µM), indicative of even higher affinity. Additionally, ComGC was enriched from extracts of cells grown to competence via ssDNA affinity together with known ssDNA binding proteins, and with periplasmic DNA receptor ComEA, which has also been found to bind to ssDNA, albeit with lower affinity than to dsDNA (37). These data establish that *Bacillus* competence pili are ds-as well as ssDNA binding machineries. Besides minor pilin ComGD, ComGF was also found in our pull-down experiments, indicating that this predicted minor pilin may also contribute to ssDNA binding. In accordance with current *in vivo* studies of *Streptococcus sanguinis* and *Streptococcus pneumoniae* (55,59), we were able to identify a residue in ComGD that are involved in DNA binding *in vitro* and *in vivo.* In contrast to other organisms, *Bacillus* seems to bind both single and double stranded DNA through minor pilins as well as major pilin ComGC. In contrast to e.g. *Streptococcus* species for which it was reported that mutations in minor pilins completely eliminate DNA binding and uptake (54,55), lowered ssDNA binding capacity in *B. subtilis* ComGD lowered transformation efficiency markedly, but not dramatically, in agreement with the finding that ComGC also binds to ssDNA.

Efficient binding of ssDNA opens the possibility to bind to single stranded regions likely existing in eDNA; binding to internal ssDNA regions could lead to pulling of this flexible part through the cell wall, followed by further pulling of the remaining dsDNA regions (Fig. 8). Complete uptake into the periplasm could be achieved by ComEA, analogous to the ratchet mechanism shown to operate in *V. cholerae* (69). In the *B. subtilis* natural habitat, the soil, a large extent of environmental DNA (eDNA) is present, with concentrations ranging from 0.03–200 μg DNA/g soil (70). Even higher concentrations of eDNA have been measured in the supernatant for undomesticated *B. subtilis* strain 3610, at the end of exponential growth phase, with 2.4–6 µg ml^−1^ eDNA (71). This high abundance of eDNA might be predominantly used for HGT-i.e. associated with chromosomal integration of DNA or may serve as a nutrient source. E-DNA can be divided into two prevailing species, namely ssDNA and dsDNA (72), and evidence has been put forward that ssDNA can be defined as the dominant form of the eDNA pool (73). Alongside these two species, the presence of hybrid DNA (i.e. DNA a composed of single and double stranded stretches) is highly conceivable.

**Figure 8:**
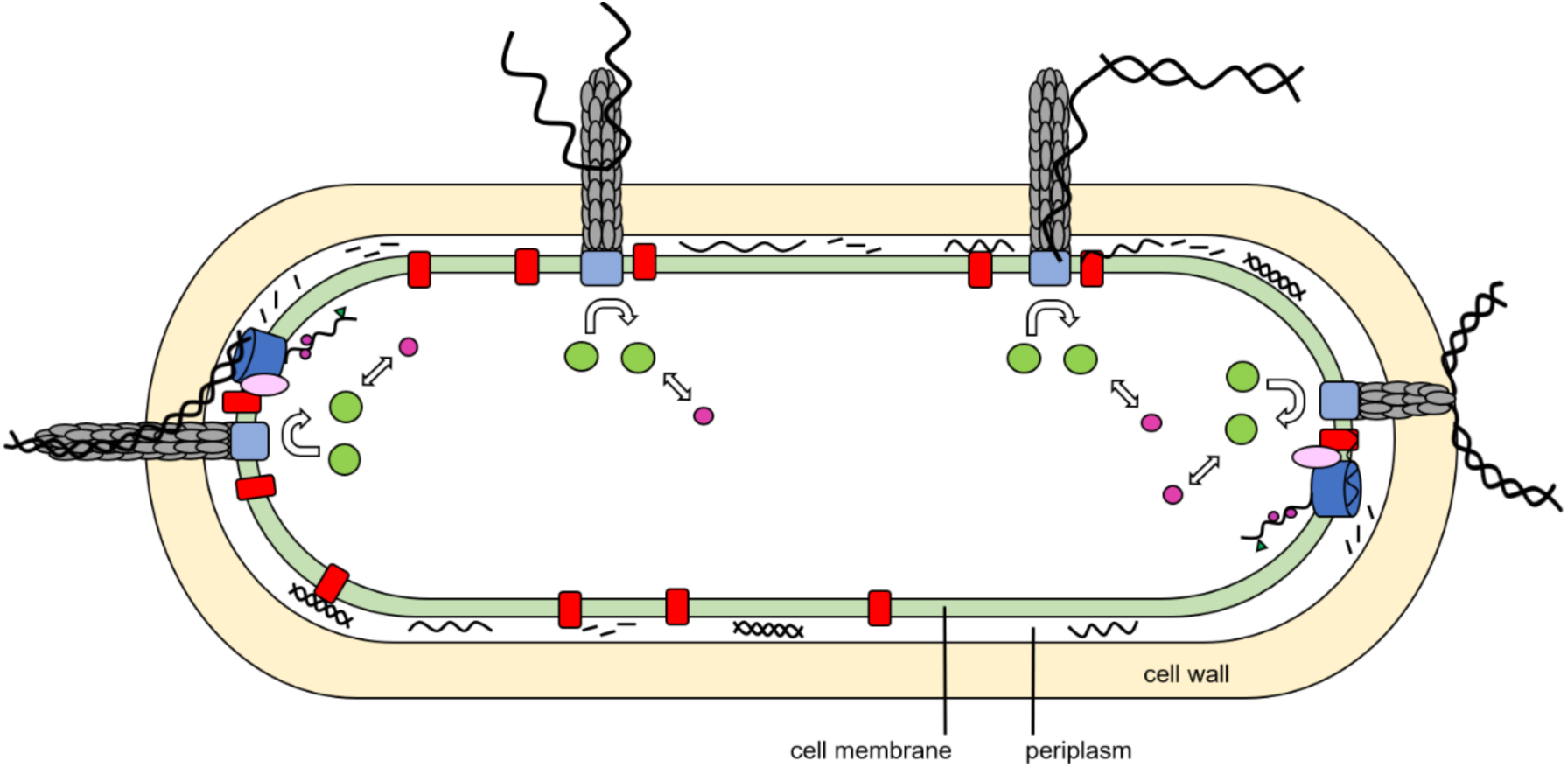
Model for ssDNA and dsDNA uptake in competent *B*. *subtilis* cells. Pilus structures retract to pull ssDNA or dsDNA inside competent *B. subtilis* cells, where ComEA will receive the DNA. Pili consist of minor and major pilins that bind to DNA, a putative platform protein (light blue) and ATPase ComGA, which may provide energy for assembly. DNA will diffuse within the periplasm alone or bound to ComEA. Within the periplasm, endonuclease NucA stochastically cuts dsDNA, apparently to facilitate uptake through the cell membrane (64). At the poles, DNA will be further taken up into the cytosol by ComEC (dark blue) and the ATPase ComFA (light purple). Inside the cytosol ssDNA binding proteins (purple circle) and DprA (green triangle) will bind to the DNA which can be further integrated into the chromosome.

Interestingly, by tracking single ssDNA molecules in real time, we found a comparable distribution of diffusion constants for ssDNA as have been reported for dsDNA (46), although the tested dsDNA molecules were of much greater length than ssDNA oligonucleotides. Possibly, dsDNA molecules are degraded into short pieces of DNA, with long stretches (required to achieve homologous recombination) being an exception rather than the rule. In any event, taken up ssDNA appears to behave like dsDNA in terms of molecule dynamics within the periplasm, and is clearly efficiently transported like dsDNA.

Previous studies using single molecule force measurements *in vivo* led to the proposal of a two-step binding for DNA pilin interaction, where an initial binding step is of transient nature and a second more rare step leads to the stable formation of the DNA/protein hybrid with efficient uptake of several tens of thousands of base pairs (74). Our quantification of DNA-pilin interaction would support this model as the K_D_ values in the micromolar range that were determined would lead to frequent transient binding events, which become highly stable when the captured DNA molecule interacts with an increasing number of pilis along the pilus. It is also compatible with our suggestion that for uptake of dsDNA, binding of DNA ends are required, while binding of the pilus tip internally to a dsDNA strand would not result in efficient transport through the cell wall.

Different models have been proposed for the biological meaning of DNA uptake, including the idea that the DNA serves as a nutrient source, or uptake of environmental DNA is employed for DNA repair mechanisms (75). In support of this idea, YhaM protein involved in the uptake of extracellular DNA was shown to be under the control of an SOS-induced promoter (63). In any event, high affinity ssDNA binding by competence pili allow cells to acquire carbon, nitrogen and phosphorous sources from soil, possibly with much higher efficiency that by bacteria lacking such structures, which depend on exoenzymes making nucleotides available from eDNA as a common good. In *N. gonnorhoae,* the uptake of environmental ssDNA predominantly serves as a source of nutrients due to its degradation in competent cells (76). Possibly, this ability is much more widespread in bacteria than anticipated before.

Membrane permease ComEC contains an OB-fold domain, which can bind to ss - as well as dsDNA; the nuclease domain can degrade one strand from dsDNA, but also ssDNA when it is bound at its 5’ end; ssDNA binding is even stronger than binding of dsDNA (77). Thus, it appears that as ssDNA can be efficiently imported into the *B. subtilis* periplasm, it should also be able to serve as transforming DNA. To prove this, long ssDNA containing a resistance cassette (usually 1500 nt) plus 500 nt flanking regions would be required, wherefore the idea that ssDNA can efficiently be used as transforming DNA in *B. subtilis* must still await confirmation.

In conclusion, we propose an important extension of the current model of DNA uptake via dynamic pili, which is conserved among Gram-positive and Gram-negative bacteria, with the addition of hybrid and single stranded DNA as efficient substrates. According to the model, taken up DNA forms first accumulates in the periplasm in all species, largely bound by ComEA, followed by either conversion of dsDNA into ssDNA, or direct transport of ssDNA by the ComEC channel, which is located at a single cell pole in *B. subtilis* (Fig. 8). Incoming cytosolic ssDNA is the substrate for RecA-dependent recombination and import of this species prevents uptake of virus dsDNA. In the absence of any homology, taken up DNA is converted into nucleotides available for scavenging (4).

## Acknowledgements

We would like to thank Dr. Rebecca Hinrichs for help on the manuscript, Lucas Schnabel, Sarah Jakob, Dr. Uwe Linne and Dr. Wieland Steinchen and Patrik Brück for technical advice. Additionally, we would like to thank Prof. Dr. Gert Bange and Prof Dr. Martin Thanbichler for technical support. This work was supported by funding from the Deutsche Forschungsgemeinschaft.

