## SUPPLEMENTARY DATA for "*Bacillus* competence pili are efficient single - and double stranded DNA uptake machines"

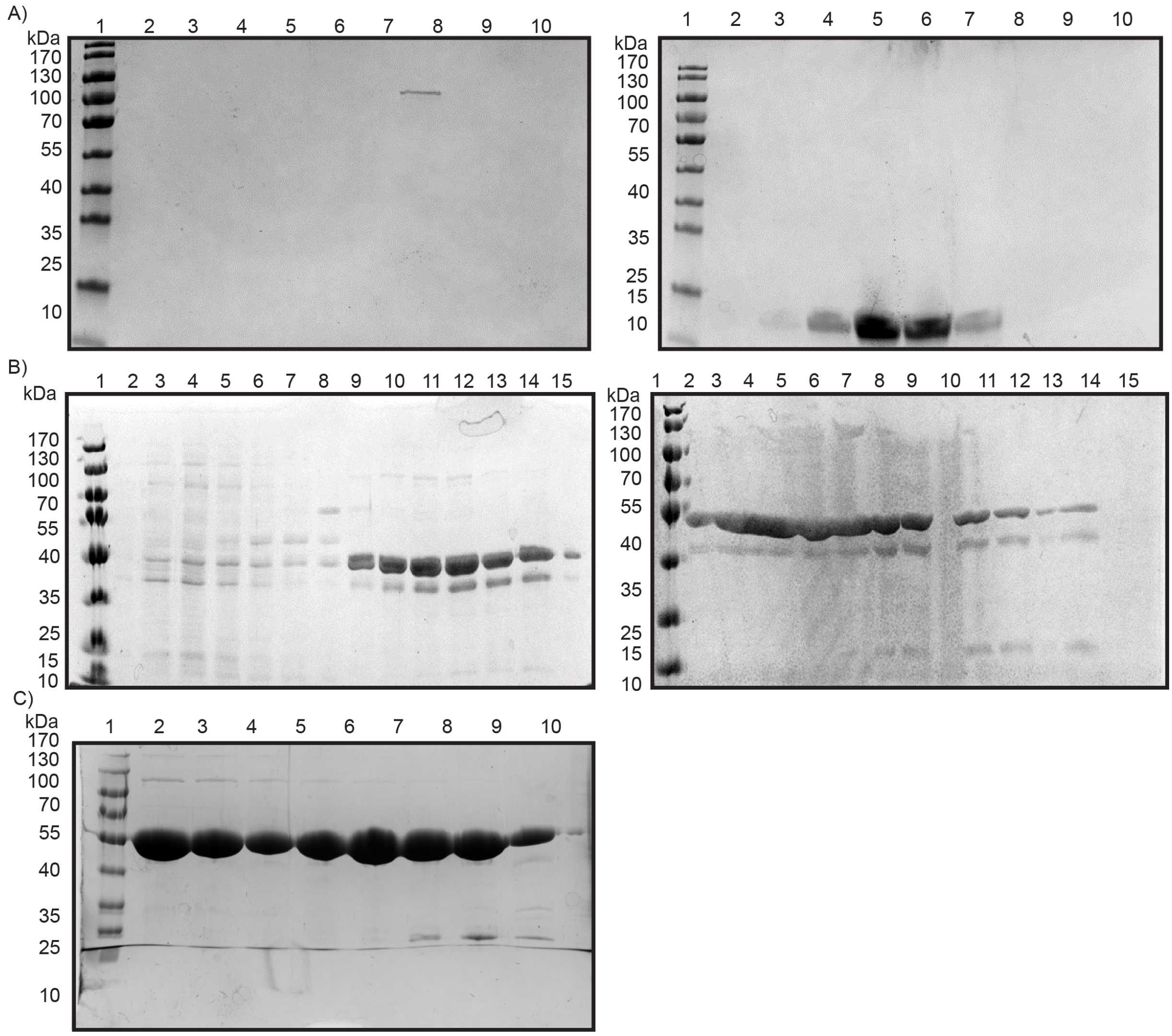


**Figure S1:** SDS PAGE (containing denaturing and reducing agents) of size exclusion runs of *Bs*ComGC, *Gt*ComGC and *Gt*ComGD. A) Preparative size exclusion of *Bs*ComGC. Left Coomassie stained SDS PAA gel represents void fractions (2-10) and right gel of monomeric fractions (2-10) corresponding to ComGC at around ~10 kDa. Note that the right panel is also shown as Fig. 1G. Line 1 in left and right gel represents PageRuler Prestained Protein Ladder. Elution profiles of size exclusion runs are shown in figure 1E. B) Preparative size exclusion run of *Gt*ComGC with void fractions (2-8), dimeric (9-15 left gel; 2-7; right gel) and monomeric fractions (8-14 right gel, no sample loaded at line 10). Lane 1 represents Page-ruler. Elution profiles of size exclusion runs are shown in main text figure 2. C) Preparative size exclusion run of *Gt*ComGD of dimeric (lane 2-7), and monomeric fractions (8-10). Line 1 in each SDS PAA gel represents PageRuler Prestained Protein Ladder. Elution profiles of runs are shown in figure 3.

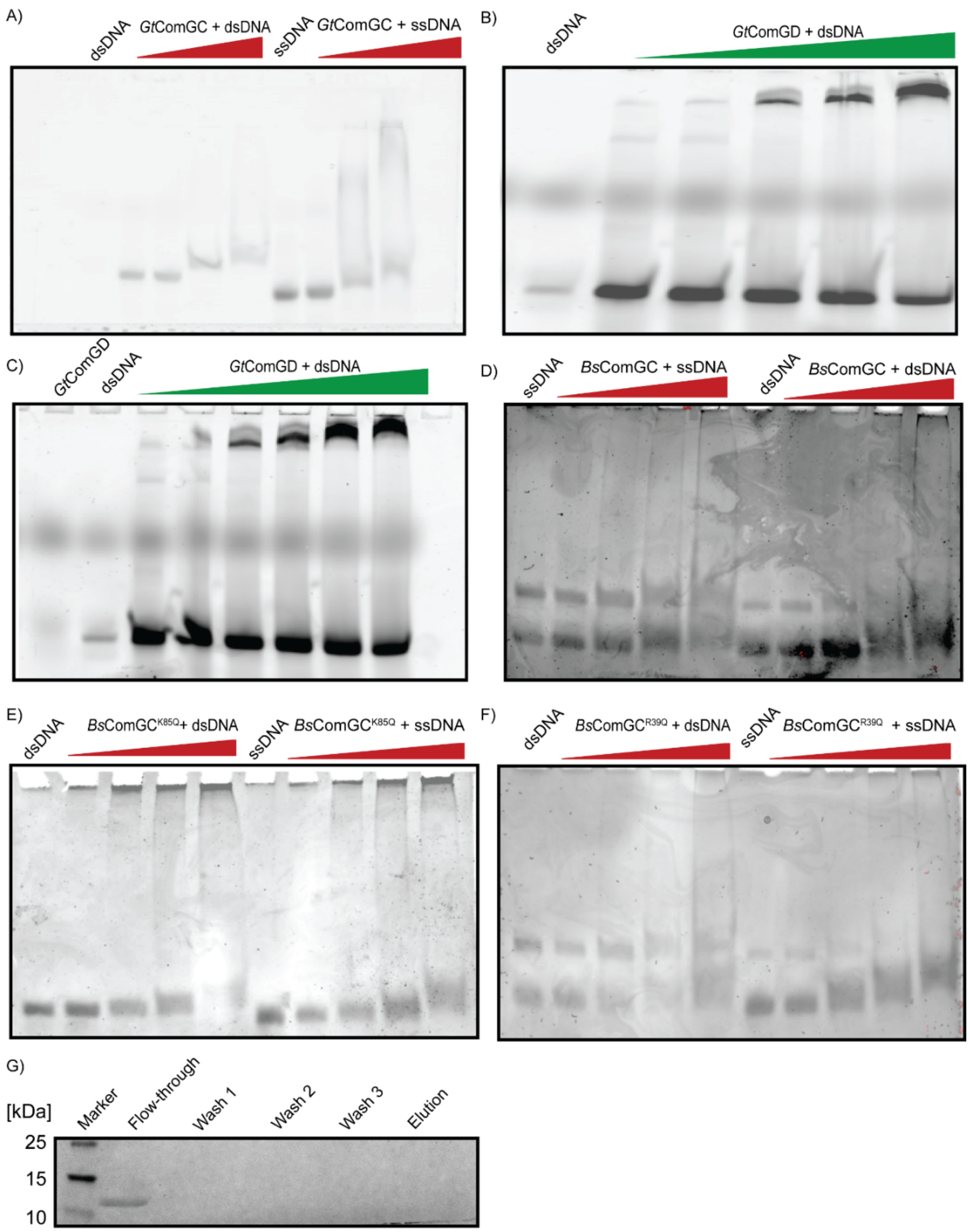


**Figure S2: DNA binding properties of major pilin *Gt*ComGC, *Bs*ComGC as well as *Gt*ComGD.** A) Native page of electromobility shift assays of major pilin of *B. subtilis* and *G. thermodenitrificans* including protein variants with A) 50 nt partially single-stranded (Fig. S3E) or 50 bp dsDNA with *Gt*ComGC (concentrations 0/50/100/150 µM). B) and C) of HIS-MBP-ComGD of mono- as well as dimeric fractions. D) 50 nt partially single - or dsDNA with major pilin *Bs*ComGC and E) variants of *Bs*ComGC$^{K85Q}$ and F) of *Bs*ComGCR$^{39Q}$. Primer sequences in table S1. DNA was incubated with increasing concentrations of protein from 20 to 50 µM. For visualization DNA was either detected by a typhoon scanner (A, B, C) or stained by Midoori green and detected by UV-light (D, E, F). G) Coomassie stained SDS PAGE, control immobilization experiments using *Bs*ComGC with Streptactin-agarose beads (lacking bound DNA) incubated with competent *B. subtilis* lysate. Lanes represent 1) PageRuler Prestained Protein Ladder, 2) Flow through, 3-5) wash steps and 6) elution fractions.

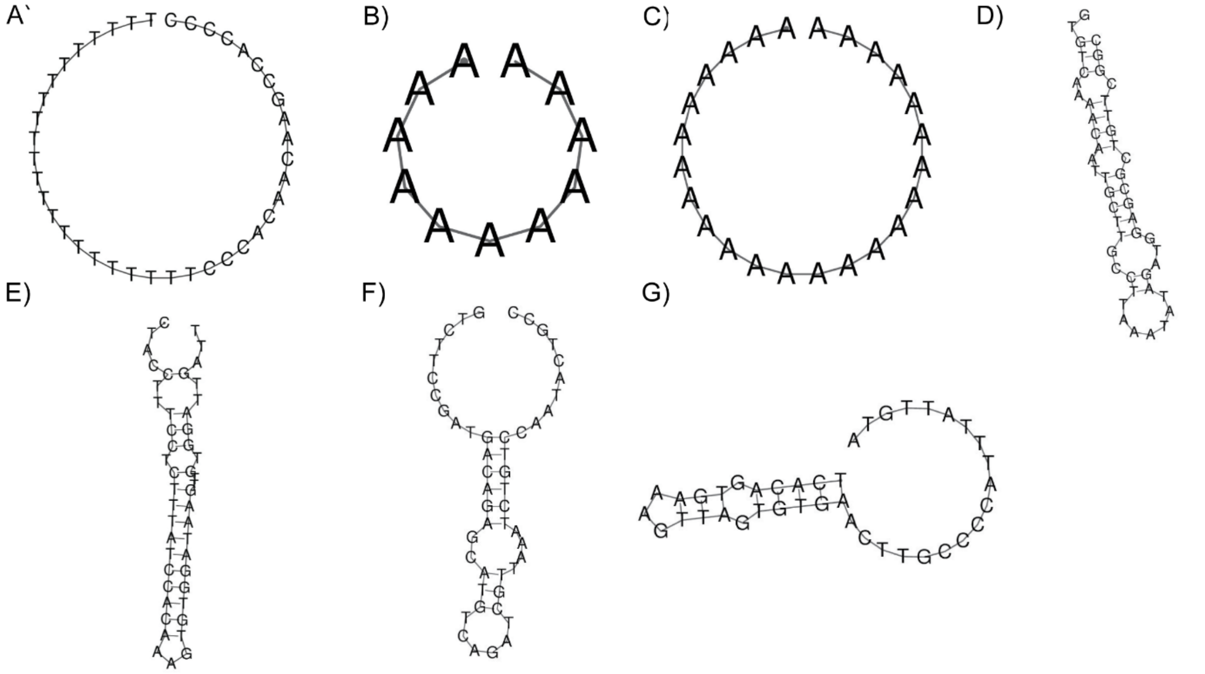


**Figure S3:** Prediction of secondary structures of used ssDNA. A-G) SsDNA was used for EMSA experiments, biolayer interferometry analysis, immobilization experiments and Single molecule tracking. Formation of secondary structures was predicted by vector builder (DNA Secondary Structure Prediction Tool | VectorBuilder). Modification, sequence and usage are listed in table S1.

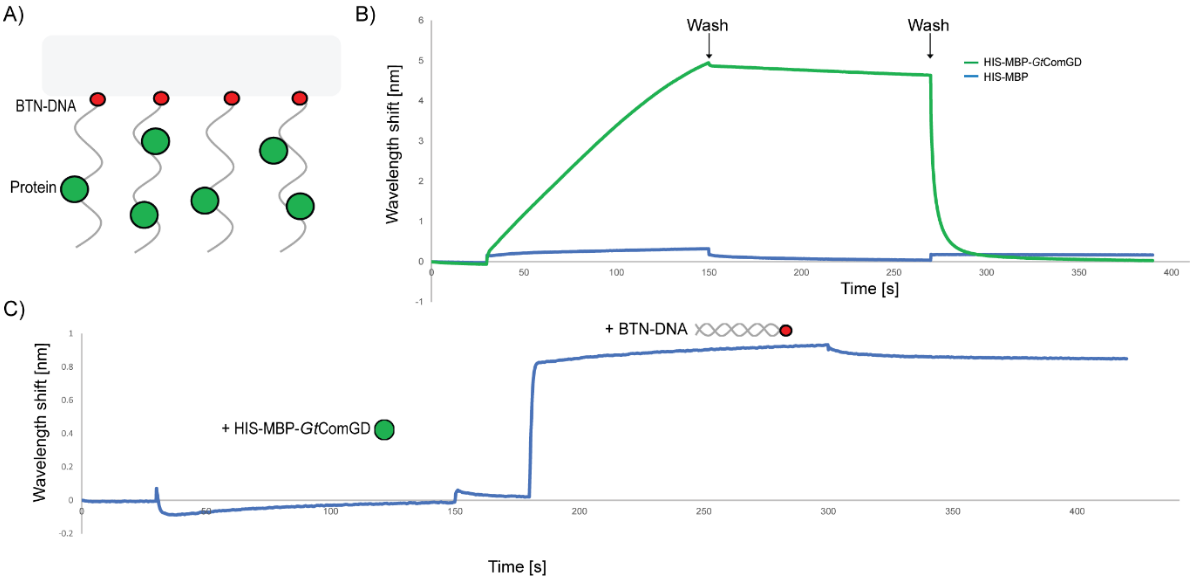


**Figure S4:** Biolayer interferometry analysis to test DNA binding. A) Schematic illustration of experimental set up. Biotinylated double- or single stranded DNA was immobilized on a SAX2 biosensor. Increasing protein concentrations were added and protein binding was followed by the detection of a concentration dependent wavelength shift e.g. shown in B). To rule out unspecific binding of the tag to the DNA loaded biosensor, His-MBP (blue line, 40 µM) was loaded in comparison to His-MBP-ComGD (green line, 28 µM). Washing steps are indicated with (Wash 1, low salt concentration and Wash 2, high salt concentrations). C) To additionally exclude the possibility that the protein itself binds to the biosensor, the protein was tested without prior loading of DNA (indicated as + His-MBP-*Gt*ComGD) in comparison to biotinylated ssDNA (indicated as +BTN-DNA). In between the loading steps (protein or DNA), baselines are shown.

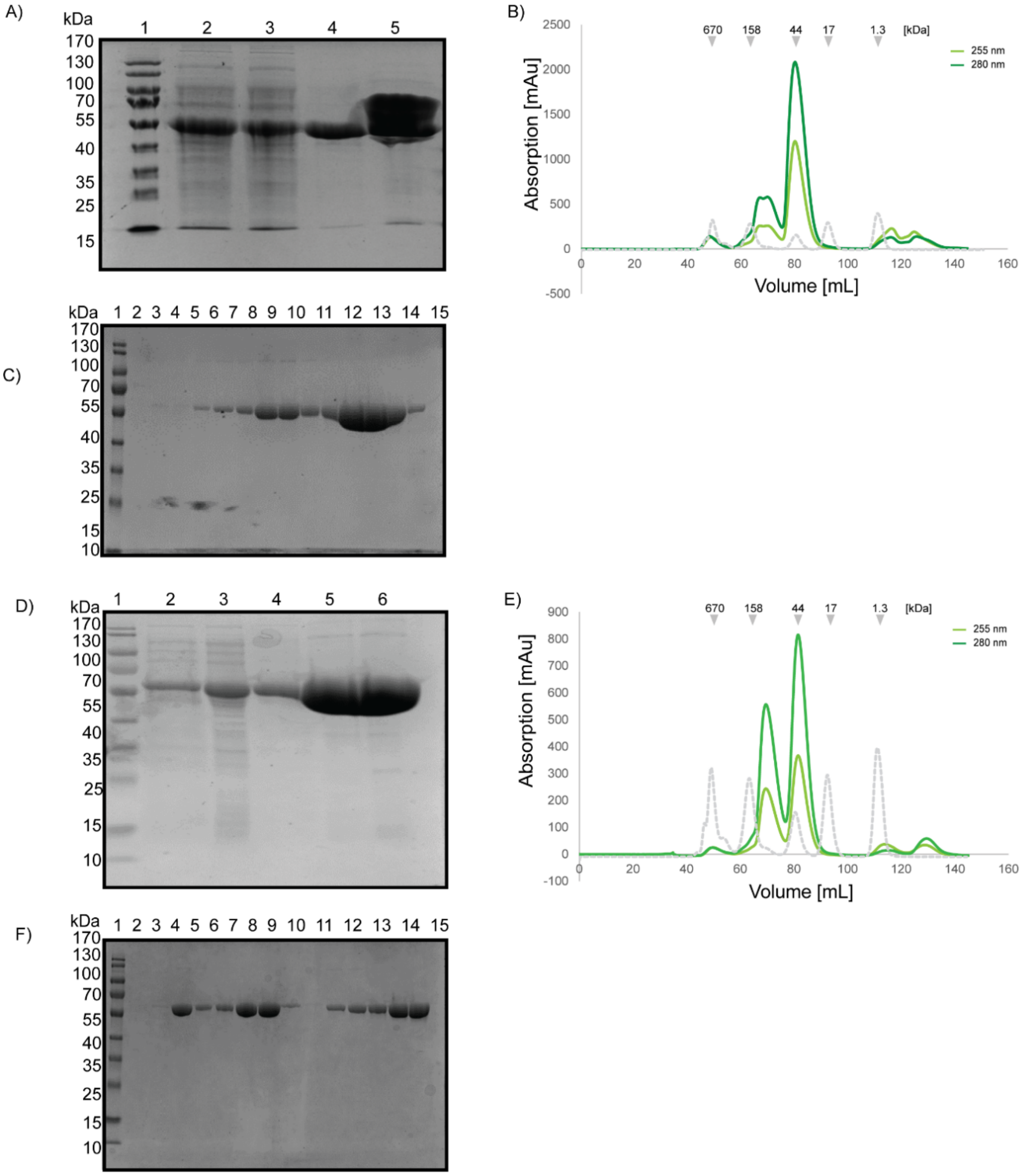


**Figure S5:** Purification of His-MBP-*Gt*ComGD variants. A) Coomassie stained SDS PAA gel of Ni-NTA purification steps of His-MBP-*Gt*ComGD$^{R139Q}$ by Ni-NTA chromatography. Lane 1 represents PageRuler Prestained Protein Ladder, lane 2) lysate, 3) flow through, 4) wash and 5) elution fraction. B) Preparative size exclusion. Light green represents absorption at 255 nm and dark green at 280 nm. C) Coomassie stained SDS PAA gel of size exclusion run of His-MBP-*Gt*ComGD variant, Lane 2-4 represent void fractions and 5-15 dimeric and monomeric fractions. D) Coomassie stained SDS PAA gel of purification of Ni-NTA purification steps of *Gt*ComGD$^{K137Q}$ (Lanes 8-12). E) Preparative size exclusion runs of His-MBP-*Gt*ComGD$^{K137Q}$. Light green shows absorption at 255 nm, dark green at 280 nm. Coomassie stained SDS-PAGE is shown in F). Lane 2-3 represents void fractions, and 4-15 dimeric and monomeric fractions. If not stated otherwise lane 1 represents PageRuler Prestained Protein Ladder.

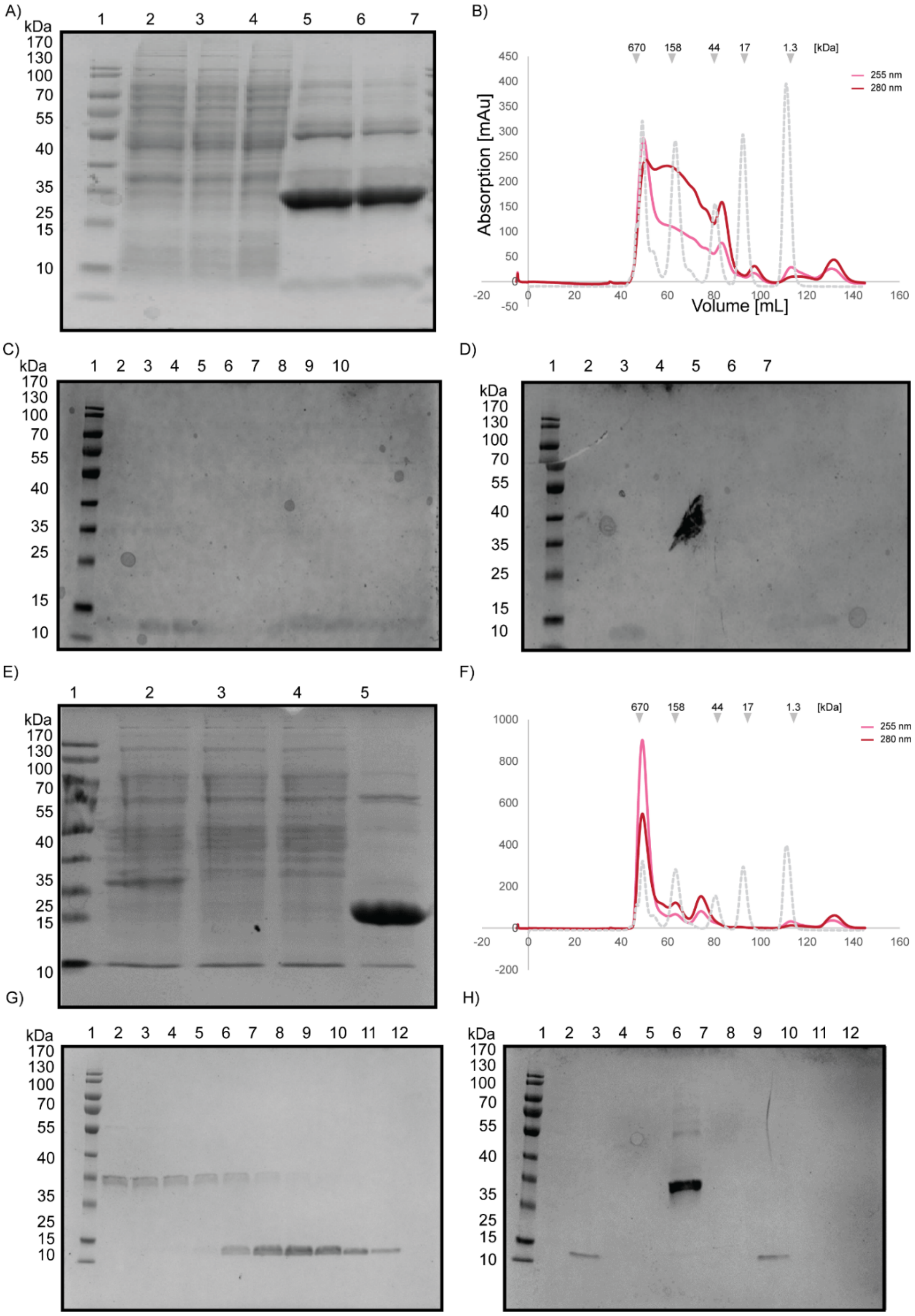


**Figure S6:** Purification of *Bs*ComGC variants. A) Coomassie stained SDS PAA gel of Ni-NTA purification steps of HIS-GST-*Bs*ComGC$^{K39Q}$. Lane 1 represents PageRuler Prestained Protein Ladder, lane 2) Lysate, 3) Flow through, 4) wash and 5,6) elution fraction. Before Size exclusion protein was treated with TEV protease and separated by a B) preparative size exclusion. Light red indicates signal at 255 nm and dark red at 280 nm. C) Coomassie stained SDS PAA gel of preparative size exclusion run of fig.S3B. ComGC eluted at around ~100 ml (monomeric fraction) with light signal of GST (lane 2-10) at ~30 kDa. To separate GST/ ComGC fractions a reverse Ni-NTA purification was used. Respective Coomassie stained SDS PAA gel is shown in fig D) Lanes represent 1) PageRuler Prestained Protein Ladder, 2) Flow-through, lane 3-4) Washing steps and lane 5) elution fraction. E) Coomassie stained SDS PAA gel of purification of Ni-NTA purification steps of HIS-GST-*Bs*ComGC$^{K85Q}$. Lane 2 represents lysate, lane 3) flow through, lane 4) washing steps and lane 5) elution fraction (treated with TEV protease). Before protein was treated with TEV protease impurities were removed by F) preparative size exclusion chromatography. Dark red indicates signal at 280 nm, light red at 255 nm. Tag was removed after size exclusion by a reverse Ni-NTA purification. Respective Coomassie stained SDS PAA gel is shown in G) Lane 1 represents PageRuler Prestained Protein Ladder, lane 2 Flow through, 3-5 washing steps and lane 6 elution. Protein was found in washing fractions. H) Comasse stained SDS PAA gel of reverse-nickel NTA with lane 1) PageRuler Prestained Protein Ladder and 2-8 respective flow through, washing and elution fractions (8).

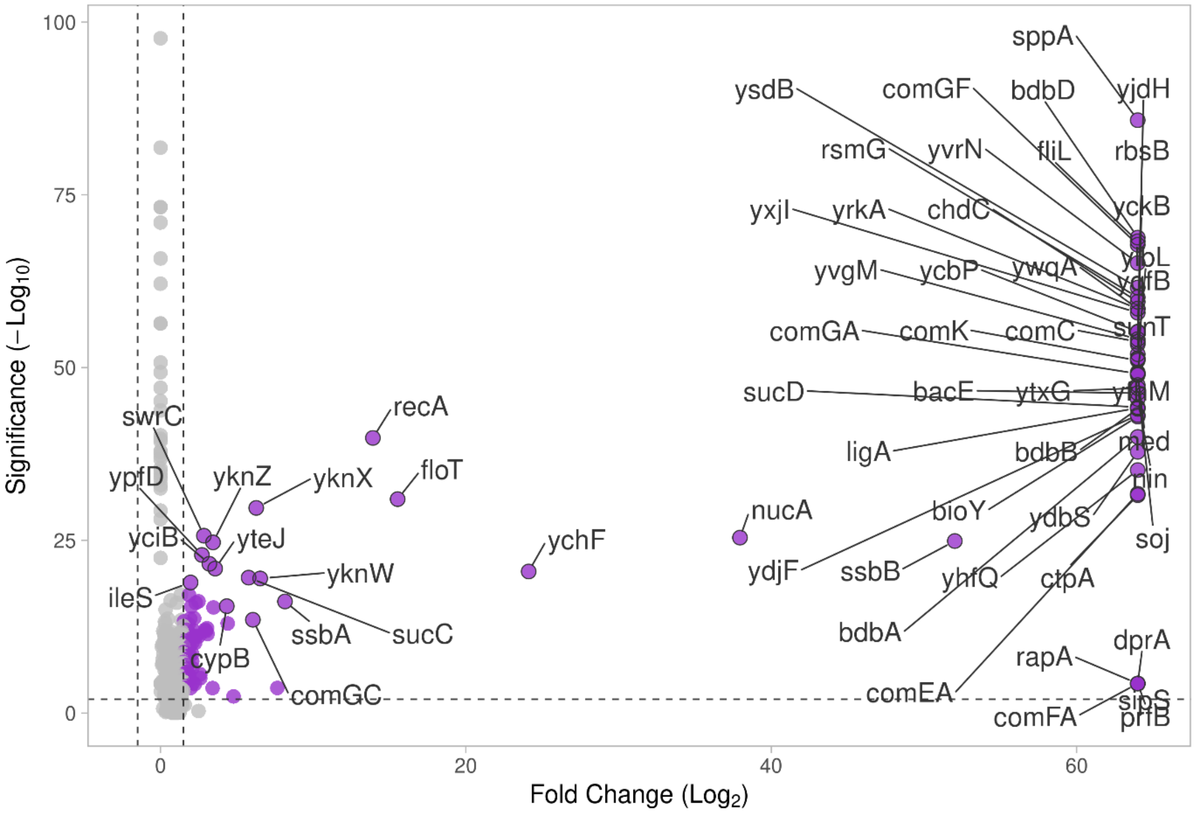


**Figure S7:** LFQ analysis of immobilization experiment with immobilized ssDNA and bound proteins of lysates from a Δ*rok* strain compared with a Δ*comK* strain. Data was blotted with VolcanoseR. Name of genes indicate bound proteins that were identified based on a significant difference compared to the negative (Δ*comK*) control.

.

**Table S1:** Unique peptides of selected proteins of immobilization experiments which were bound to single/ or double stranded DNA (of *B. subtilis* lysate) determined by Mass spectrometry. Replicates are indicated with numbers (1-9). In 1-3 lysate of Δ*comK* strain was used, from 4-9 lysate of PY79, Δ*rok* was used.

| **Protein** | **1** | **2** | **3** | **4** | **5** | **6** | **7** | **8** | **9** |
|---|---|---|---|---|---|---|---|---|---|
| Single-stranded DNA-binding protein A GN=*ssbA* | 19 | 4 | 10 | 2 | 15 | 31 | 7 | 26 | 7 |
| ComG operon protein 3 GN=*comGC* | 0 | 0 | 0 | 3 | 4 | 5 | 2 | 3 | 6 |
| ComE operon protein 1 GN=*comEA* | 0 | 0 | 0 | 32 | 58 | 25 | 45 | 26 | 52 |
| Single-stranded DNA-binding protein A GN=*ssbB* | 0 | 0 | 0 | 9 | 6 | 13 | 10 | 16 | 13 |

**Table S2** Statistics of labelled ss/dsDNA with various lenths.

| | **ssDNA [25 bp]** | **ssDNA [40 bp]** | **Hairpin-DNA [48 bp]** |
|---|---|---|---|
| Cells | 182 | 149 | 65 |
| Cells with tracks | 68 | 114 | 43 |
| Total cells/Cells with tracks [%] | 37 | 76 | 67 |
| tracks | 115 | 447 | 365 |
| Tracks/ cell | 1.49 | 3.79 | 6.73 |

**Table S3** Statistical analysis (SQD) analysis of labelled ssDNA with various lengths using simultaneous fitting.

| | ssDNA [25 bp] | ssDNA [40 bp] | hairpin-DNA [48 bp] |
|---|---|---|---|
| **$Pop_1$ [%]** | 22.8 +/- 0.005 | 29.9 +/- 0.005 | 23.5 +/- 0.005 |
| **$Pop_2$ [%]** | 54 +/- 0.005 | 46.6 +/- 0.005 | 46.7 +/- 0.005 |
| **$Pop_3$ [%]** | 23.2 +/- 0.005 | 23.5 +/- 0.01 | 29.7 +/- 0.022 |
| **$D_1$ [$\mu m^2$ $s^{-1}$]** | 0.009 +/- 0 | 0.009 +/- 0 | 0.009 +/- 0 |
| **$D_2$ [$\mu m^2$ $s^{-1}$]** | 0.06 +/- 0.001 | 0.06 +/- 0.001 | 0.06 +/- 0.001 |
| **$D_3$ [$\mu m^2$ $s^{-1}$]** | 0.51 +/- 0.01 | 0.51 +/- 0.01 | 0.51 +/- 0.01 |

**Table S4** Statistical analysis (SQD) analysis of labelled ssDNA with various lengths using individual fitting.

| | ssDNA [25 bp] | ssDNA [40 bp] | hairpin-DNA [48 bp] |
|---|---|---|---|
| **$Pop_1$ [%]** | 30.5 +/- 0.0012 | 27.9 +/- 0.002 | 27 +/- 0.003 |
| **$Pop_2$ [%]** | 45.9 +/- 0.005 | 47.3 +/- 0.002 | 45.3 +/- 0.002 |
| **$Pop_3$ [%]** | 23.6 +/- 0.005 | 24.9 +/- 0.01 | 27.7 +/- 0.002 |
| **$D_1$ [$\mu m^2$ $s^{-1}$]** | 0.012 +/- 0 | 0.008 +/- 0 | 0.002 +/- 0 |
| **$D_2$ [$\mu m^2$ $s^{-1}$]** | 0.07 +/- 0.004 | 0.07 +/- 0.001 | 0.07 +/- 0.001 |
| **$D_3$ [$\mu m^2$ $s^{-1}$]** | 0.46 +/- 0.01 | 0.47 +/- 0.01 | 0.57 +/- 0.01 |

**Table S5: Bacterial strains used in this study.**

| Strain | Genotype | Usage | Reference |
|---|---|---|---|
| DH5α | F⁻ *endA1 glnV4 4 thi1 recA1 relA 1 gyrA96 deoR nupG purB20* φ 80d*lacZ*ΔM15 Δ(*lacZYA-argF*)U169, hsdR17($r_K^-m_K^+$), λ⁻ | Cloning | Thermo scientific™ |
| C41 (DE3) | F- ompT hsd SB (r- mB-) gal dcm (DE3) | Heterologous expression | Sigma-Aldrich (1) |
| PY79 | Laboratory strain | Transformation efficiency experiments of *B. subtilis* | |
| Δ*rok* | PY79, Δ*rok* | Pull-down experiments | (2) |
| Δ*rok*, Δ*comEC* | PY79, Δ*rok*, Δ*comEC* | Single-molecule tracking | (3) |
| Δ*comk* | PY79, Δ*comK* | Pull-down experiments | (4) |
| ComGD$^{\mathbf{R137Q}}$ | PY79, ComGDR$^{137Q}$ | Transformation efficiency experiments of *B. subtilis* | This work |

**Table S6:** Oligonucleotides used in this study. Oligonucleotides were used for molecular cloning, Crispr-cas9, DNA-binding experiments (immobilization assays, Biolayer interferometry and Electromobility shift assays), single-molecule tracking and *in vivo* analysis by transformation efficiencies of *B. subtilis* and *H. pylori.*

| **Name** | **sequence** | **Usage** |
|---|---|---|
| BS_ComGC 28 FWD | GAGGGTCTCCCATGGGCAACGTCACGAAA CATAATCAAACCATTC | Molecular cloning construct for *Bs*ComGC truncation |
| *BS*_ComGC REV | AGGAGGGTCTCCTCGAGTCTTAATGTTCAA CCTTAACTTCTCCGCC | Molecular cloning construct for *Bs*ComGC truncation |
| *GT*_ComGD 29 FWD | GAGGGTCTCCCATGGGCGAGCTTGGTGGG ATCATGCAGC | Molecular cloning construct for *Gt*ComGD truncation |
| *GT*_ComGD 29 REV | AGGAGGGTCTCCTCGAGTTACATTTTTTGT ACATAAAACCGCCCTTTTCCG | Molecular cloning construct for *Gt*ComGD truncation |
| *GT*_ComGC 28 FWD | GAGGGTCTCCCATGGGCAACATCACAAAG CATAACGGCATG | Molecular cloning construct for *Gt*ComGC truncation |
| *GT*_ComGC 28 REV | AGGAGGGTCTCCTCGAGTTACGAGCCAAT TTCGCTCACATCG | Molecular cloning construct for *Gt*ComGC truncation |
| *BS* ComGC $^{K85Q}$ FWD | TCCAAATGGTCAGCGCATTATCATC | Point mutation in ComGC |
| ComGC$^{K39Q}$ REV | GTTTGATTATGTTTCGTGAC | Point mutation in ComGC |
| *BS* ComGC $^{K39Q}$ FWD | CATTCAAAAACAGGGCTGTGAAG | Point mutation in ComGC |
| *BS* ComGC $^{K85Q}$ REV | TCTGACTGTAAATCGGC | Point mutation in ComGC |
| *GT* ComGD $^{K137Q}$ FWD | TTTGCTCGGACAGGGGCGGTTTT | Point mutation in ComGD |
| *GT* ComGD $^{K137Q}$ REV | AACGTCACTTTATAACTGTTTTCG | Point mutation in ComGD |
| *GT* ComGD $^{R139Q}$ FWD | CGGAAAAGGGCAGTTTTATGTAC | Point mutation in ComGD |
| *GT* ComGD $^{R139Q}$ REV | AGCAAAAACGTCACTTTATAAC | Point mutation in ComGD |
| ssDNA BLi-experiments | GTG TCA AAA CAA TTG CTT GCC TTA AAT ATA GAT GGA GCG CTG TTC GGC | ssDNA for BLi-experiments [Btn] |

| | | |
|---|---|---|
| dsDNA BLi-experiments | GCTGAAACGTGTATTACATG | dsDNA for BLi experiments [Btn] |
| dsDNA BLi-experiments | CTGCCGCCACAAGTTTATAT | dsDNA for BLi experiments |
| Single molecule tracking ssDNA | CTACCTTTCCTCTTTATCCACAAAGTGTGGATAAGTTGTGGATTGATT | ssDNA for single molecule tracking [CY3] |
| ssDNA Ery FWD | GGGAACGGTTGGAGCTAATG | DsDNA EMSA |
| ssDNA/ dsDNA Ery REV | TTCCGGGAACAGTGACAGAG | dsDNA fpr EMSA |
| EMSA FWD ssDNA/ dsDNA | GTGAAAAACCCGCTCATCGATGATGGGCGGGTTTTTTTGCGGATGTTGAA | ssDNA/ dsDNA EMSA |
| EMSA REV dsDNA | TTCAACATCCGCAAAAAAACCCGCCCATCATCGATGAGCGGGTTTTTCAC | DsDNA EMSA |
| EMSA FWD ssDNA/ dsDNA | GTCTTCCGATGACAGAGCATGTCAGATCGTTAAATCTGTCCAATACTGCC | ssDNA/ dsDNA EMSA |
| EMSA REV dsDNA | GGCAGTATTGGACAGATTTAACGATCTGACATGCTCTGTCATCGGAAGAC | DsDNA EMSA |
| EMSA FWD | ATAAGGGAAGTGCAGTAAATTAGAGGAAAATCATGATTTTGTTCTCTAAAGAGAACTTATTG | ssDNA/ dsDNA EMSA, fluorescently labelled [CY5] |
| EMSA REV | CAATAAGTTCTCTTTAGAGAACAAAATCATGATTTTCCTCTAATTTACTGCACTTCCCTTAT | dsDNA EMSA |
| ssDNA Pulldown FWD | TCACAGTGAAAGTTAGTGTGAACTTGCCCCATTTATTGTA | ssDNA/ dsDNA for Pulldown [BTN] |
| dsDNA Pulldown REV | TACAATAAATGGGGCAAGTTCACACTAACTTTCACTGTGA | dsDNA for Pulldown |
| ssDNA PolyA-tail short | AAAAAAAAAAAA | Immobilization experiment [BTN] |
| ssDNA ssDNA PolyA-tail short | AAAAAAAAAAAA | Single molecule tracking [FAM] |
| ssDNA middle | AAAAAAAAAAAAAAAAAAAAAAAAA | Single molecule tracking [FAM] |
| ssDNA long | TTTTTTTTTTTTTTTTTTTTTTCCCACAACAAGCCACCCC | Single molecule tracking [FAM] |
| ssDNA long | TTTTTTTTTTTTTTTTTTTTTTCCCACAACAAGCCACCCC | Single molecule tracking [BTN] |
| gDNA comGD R137Q rev | gtatcatgaatgtaataagaaacccg | Crispr-cas9 |

| | | |
|---|---|---|
| gDNA test comGD R137Q for | acggctaaaagccggtg | Crispr-cas9 |
| pPB105 TEMP DO comGD R137Q for | gggCAGgtcaatgtCgagagaaaataaagg | Crispr-cas9 |
| pPB105 TEMP DO comGD R137Q rev | GCATAACCAAGCCTATGCCTACAGCatagaa attgctgcgtaccctcc | Crispr-cas9 |
| pPB105 TEMP UP comGD R137Q for | GCATGCTGAATTCGTAATGAGGTTCttgaatcc gctatttgtggaagc | Crispr-cas9 |
| pPB105 TEMP UP comGD R137Q rev | CTCTCGACATTGACCTGCCCGCTCCCTAG | Crispr-cas9 |
| SEQ comGD R137Q for | acacagcagactgctatttcc | Crispr-cas9 |
| Spacer 1 | AAACCAGTTTATCTAGGGAGCGGGAGAGT CAATGG | Crispr-cas9 |
| Spacer 2 | AAAACCATTGACTCTCCCGCTCCCTAGATA AACTG | Crispr-cas9 |
| oPEB218 VPCR pPB41 for | GAATGGCGATTTTCGTTCGTGAATAC | Crispr-cas9 |
| oPEB253 Seq CRISPR insert | GAAGGGTAGTCCAGAAGATAACGA | Crispr-cas9 |
| oPEB234V2 Cas9 rev | GTATTCACGAACGAAAATCGCCATTC | Crispr-cas9 |
| oPEB227 Seq Template5 | CCGTCAATTGTCTGATTCGTTA | Crispr-cas9 |
| oPEB Seq Template3 | TTCTACATTTAGGCGCTGC | Crispr-cas9 |
| oPEB232 Cas9 rev | GTATTCACGAACGAAAATCGCCATTCCTAG CAGCACGCCATAGTGACTG | Crispr-cas9 |
| oPEB232 Cas9 for | GCTGTAGGCATAGGCTTGGTTATG | Crispr-cas9 |
| oPEB218 VPCR pPB41 rev | GAACCTCATTACGAATTCAGCATGC | Crispr-cas9 |
| oPEB234V2 Cas9 rev | GTATTCACGAACGAAAATCGCCATTC | Crispr-cas9 |

**Table S7: Plasmids used in this study.**

| Plasmid | Usage | Reference |
|---|---|---|
| HIS-MBP | Protein overexpression with N-terminal HIS-MBP-tag | (5,6) |
| HIS-GST | Protein overexpression with N-terminal HIS-GST-tag | (5,7) |
| HIS-MBP-*Gt*ComGD | Protein overexpression with N-terminal HIS-MBP-tag *Gt*ComGD | This work |
| HIS-MBP-GtComGD$^{\mathbf{R139Q}}$ | Protein overexpression with N-terminal HIS-MBP-tag for *Gt*ComGD$^{R139Q}$ | This work |
| HIS-MBP-GtComGD$^{\mathbf{K137Q}}$ | Protein overexpression with N-terminal HIS-MBP-tag for *Gt*ComGD$^{R137Q}$ | This work |
| HIS-GST-*Bs*ComGC | Protein overexpression with N-terminal HIS-GST-tag for *Bs*ComGC | This work |
| HIS-GST-BsComGC$^{K85Q}$ | Protein overexpression with N-terminal HIS-GST-tag for *Bs*ComGC$^{K85Q}$ | This work |
| HIS-GST-*Bs*ComGCK$^{39Q}$ | Protein overexpression with N-terminal HIS-GST-tag for *Bs*ComGCK$^{39Q}$ | This work |
| HIS-MBP-*Gt*ComGC | Protein overexpression with N-terminal HIS-GST-tag for *Gt*ComGC | This work |
| pDG1664 | Template for amplification of DNA fragments for transformation efficiency experiments | (8) |
| oPEB232 | Crispr-Cas9, *in vivo* mutation in ComGD | (9) |
| oPEB232 | Crispr-Cas9, *in vivo* mutation in ComGD | (9) |
